# Degenerated CRISPRs widely regulate Cas expression to balance immunity and cost

**DOI:** 10.1101/2023.03.10.532045

**Authors:** Chao Liu, Rui Wang, Jie Li, Feiyue Cheng, Xian Shu, Huiwei Zhao, Qiong Xue, Haiying Yu, Aici Wu, Lingyun Wang, Sushu Hu, Yihan Zhang, Jun Yang, Hua Xiang, Ming Li

**Affiliations:** CAS Key Laboratory of Microbial Physiological and Metabolic Engineering, State Key Laboratory of Microbial Resources, Institute of Microbiology, Chinese Academy of Sciences, Beijing, China; State Key Laboratory of Microbial Resources, Institute of Microbiology, Chinese Academy of Sciences, Beijing, China; College of Life Science, University of Chinese Academy of Sciences, Beijing, China; College of Plant Protection, Shandong Agricultural University, Taian, Shandong, China; School of life Sciences, Hebei University, Baoding, Hebei, China; Center for Life Science, School of Life Sciences, Yunnan University, Kunming, China

## Abstract

CRISPR RNAs (crRNAs) and Cas proteins together provide prokaryotes with adaptive immunity against genetic invaders. How Cas expression is fine-tuned to avoid energy burden while satisfying the dynamic need of crRNAs remains poorly understood. Here we experimentally demonstrated widespread degenerated mini-CRISPRs encode CreR (<u>C</u>as-<u>re</u>gulating) <u>R</u>NAs to mediate autorepression of type I-B, I-E and V-A Cas proteins, based on their partial complementarity to *cas* promoters. This autorepression decreases energy burden and autoimmune risks, thus mitigating the fitness cost on host cell, and remarkably, senses and responds to alterations in the volume of canonical crRNAs, which compete with CreR for Cas proteins. Moreover, CreR-guided Cas autorepression can be subverted by diverse anti-CRISPR (Acr) proteins that destruct Cas proteins, which in turn replenishes the weapon depot. Our data unveil a general degenerated crRNA-guided autorepression paradigm for diverse Cas effectors, which highlights the intricate (self-)regulation of CRISPR-Cas and its transcriptional counterstrategy against Acr attack.

## INTRODUCTION

CRISPR-Cas systems constitute the adaptive defense line against invading genetic elements, like viruses (phages) and plasmids (Barrangou and Horvath, 2017; Brouns *et al*., 2008; Wiedenheft *et al*., 2012; Hille *et al*., 2018; Nussenzweig and Marraffini, 2020). These systems are highly diversified and currently classified into two classes, six types, and more than 30 subtypes (Makarova *et al*., 2020). CRISPR-Cas systems consist of CRISPR arrays that store invader-derived sequences spacing each two direct repeats (namely spacers), and *cas* (CRISPR-associated) genes that encode a multi-subunit effector complex (class 1) or a single-protein effector (class 2). Mature crRNAs guide the Cas effector to precisely recognize and cleave the foreign nucleic acids based on the perfect complementarity between their spacer portion and the target site (namely protospacer), wherein a conserved protospacer adjacent motif (PAM) plays a critical role during target recognition (Semenova *et al*.,2011; Wiedenheft *et al*., 2011). A typical CRISPR-Cas system usually also encodes Cas proteins (e.g., Cas1, Cas2, and Cas4) that mediate the acquisition of new spacers from the genetic invaders (Sternberg *et al*., 2016; Li *et al*., 2014; Datsenko *et al*., 2012).

CRISPR arrays are persistently acquiring new spacers and losing old ones during the conflicts between bacteria and phages, thus resulting in a dynamic volume of CRISPR memory (Savitskaya et al., 2017; Levin et al., 2013). In theory, to ensure effective CRISPR immunity, a sufficient supply of Cas proteins is required. Yet, on the other hand, their excessive production will inevitably cause energy waste and likely other deleterious effects (e.g., autoimmunity) (Stern et al., 2010; Bikard et al., 2012). Therefore, Cas expression need be precisely regulated to avoid potential fitness costs while providing sufficient protein effectors for crRNA guides. However, how Cas expression and crRNA production are coordinated remains poorly understood.

Our recent studies unraveled the regulatory function of type I-B CRISPR effector complex, which is reprogrammed by a degenerated mini-CRISPR (termed *creA* for CRISPR-resembling antitoxin) to transcriptionally repress a small toxic RNA (CreT, for CRISPR-regulated toxin) that acts by sequestering a rare tRNA species in the cell (Li et al., 2021; Cheng et al., 2021). This tiny two-RNA TA element (considered to represent type VIII TA) makes host cells addicted to CRISPR-Cas effectors, inactivation of which will liberate CreT expression and elicit cell death/dormancy. Notably, the spacer portion of CreA share limited complementarity to the promoter DNA of *creT* (P_creT_) (Figure 1A), which causes gene regulation rather than DNA cleavage. Such limited spacer-protospacer complementarity has also been reported to direct the type II effector Cas9 to transcriptionally repress a virulence-related regulon (Ratner *et al*., 2019).

**Figure 1.**
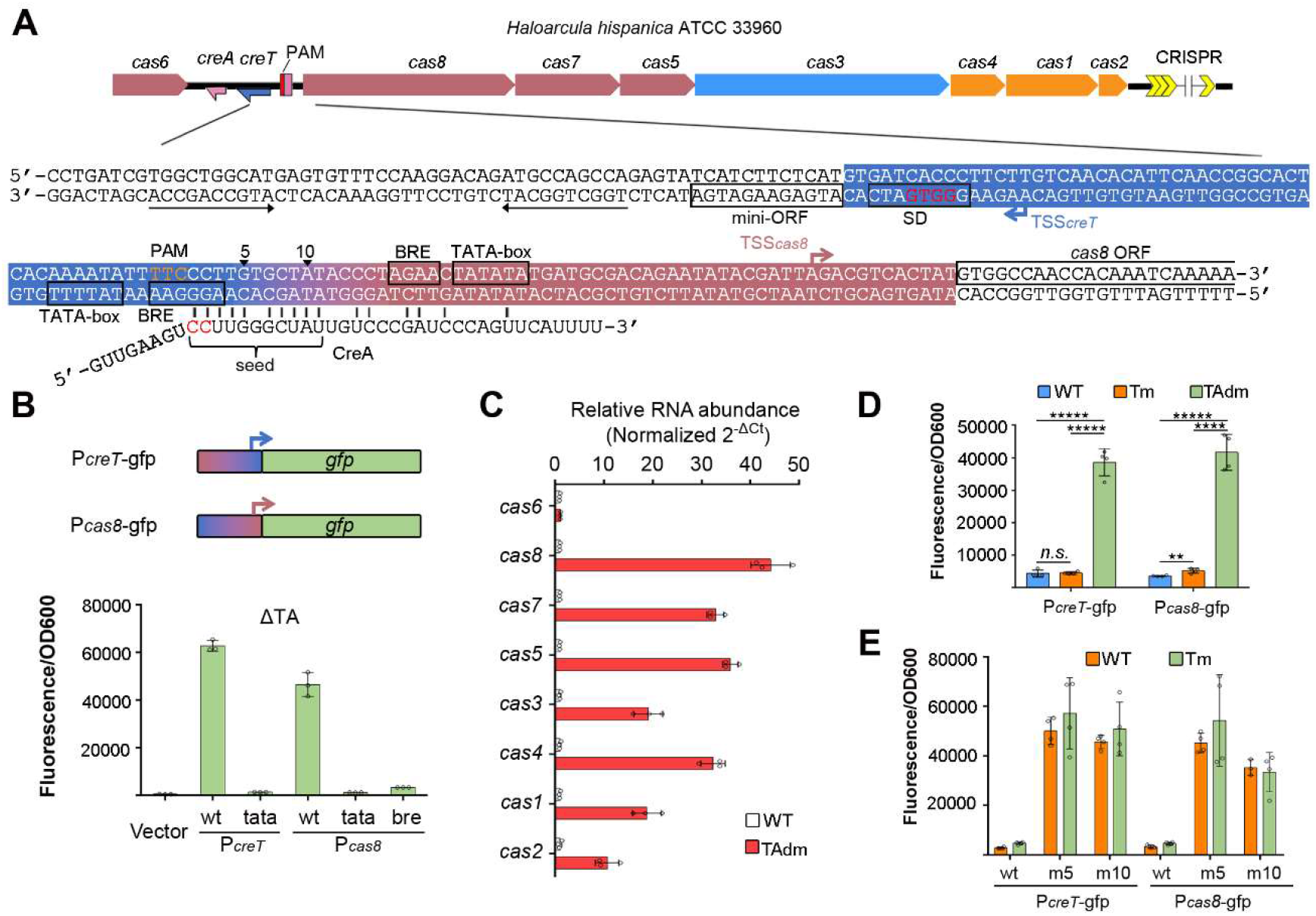
*H. hispanica* CreA synchronously represses *P_creT_* and P_cas8_. (**A**) Schematic depiction of the divergent *P_creT_* and *P_cas8_* and their targeting by CreA. Red nucleotides within the Shine-Dalgarno (SD) sequence of *creT* were mutated to generate the Tm mutant, and then red nucleotides within CreA were further mutated to construct TAdm. (**B**) Validation of *P_cas8_* using a green fluorescent protein *(gfp)* gene. The TATA-box (tata) and BRE (bre) elements were separately mutated. (**C**) The relative RNA level of *cas* genes in WT and TAdm. 7S RNA served as the internal control. (**D**) Activity of *P_creT_* and *P_cas8_* in WT, Tm, and TAdm. (**E**) Activity of *P_creT_* and *P_cas8_* in WT and Tm cells when the 5^th^ (m5) or 10^th^ (m10) seed nucleotide within the target site of CreA (labeled in panel **A**) was mutated. Error bars, mean±s.d. (n=3 or 4). Two-tailed Student’s *t* test [**P < 0.01; \*\*\*\**P* < 0.0001; *****P < 0.00001; *n.s*., not significant (*P* > 0.05)]. See also Figure S1 and S2.

In this study, we showed that CreA not only directs the type I-B Cas proteins to repress toxin expression, but also mediates their autorepression, and importantly, more creA analogs, namely CreR (for Cas-regulating RNA), were identified from type I-E and V-A systems, which also direct the autorepression circuit of their multi-subunit or single-protein effector. Remarkably, these repression circuits were relieved by elevating the expression level of canonical crRNAs, which compete for Cas proteins with these regulatory RNA guides. Hence, Cas autorepression mediated by the regulatory degenerated crRNAs (like CreA) enables the production of Cas proteins to real-time fit the volume of the canonical defensive crRNAs. In addition, we also demonstrated that this autorepression could be subverted by Acr proteins that attack Cas proteins, which illuminates a new anti-anti-CRISPR strategy that acts on transcription level.

## RESULTS

### CreA can repress both *creT* and *cas* transcription

Previously, we characterized the CreTA RNA pair that safeguards the type I-B CRISPR-Cas in the archaeon *Haloarcula hispanica* (Li et al., 2021). As this mini addiction module locates within the ~300 bp intergenic sequence between *cas6* and *cas8* (Figure 1A), we previously constructed a ΔTA mutant by simply deleting this intergenic sequence (Li et al., 2021). In a following study, we noticed that transcripts of most *cas* genes (except *cas6)* reduced by 70-90% in ΔTA (Cheng et al., 2022), which suggests a previously unnoticed promoter preceding *cas8*, referred to hereafter as *P_cas8_*. By reanalyzing the data of a small RNA sequencing assay that was originally designed to profile *creTA* transcription (Figure S1), we identified the transcription start site (TSS) of *cas8*, which locates 12 bp upstream of the open reading frame (ORF) (Figure 1A). Accordingly, we predicted the archaeal promoter elements, BRE (TF-IIB recognition) and TATA-box (Figure 1A). Using the green fluorescent protein (*gfp*) gene as a reporter, we showed that mutating either BRE or TATA-box abolished the activity of *P_cas8_* (Figure 1B). We found that *P_cas8_* was highly efficient, and was ~135 times and ~33 times the activity of *cas6* and 16S rRNA promoters, respectively (Figure S2). Surprisingly, it was also stronger than the promoter of creT (P_*creT*_; ~1.4 times) and a strong constitutive promoter we usually used for gene overexpression in haloarchaea (*P_phaR_*; ~2.0 times) (Cai *et al*., 2015).

Notably, *P_cas8_* and P_*creT*_ run divergently and tightly flank the target site of CreA (Figure 1A), suggesting their simultaneous repression by CreA. Because simply mutating CreA would lead to CreT de-repression and hence cell death, we first constructed a CreT mutant (designated as Tm) by disrupting its Shine-Dalgarno element, and then mutated the seed sequence (essential for target recognition) of CreA to generate a CreT/CreA double mutant (designated as TAdm) (Figure 1A). Surprisingly, *cas8* transcription elevated by as much as ~44-fold in TAdm (compared to WT), and the downstream *cas* genes were all markedly up-regulated (by ~10 to 36-fold); by contrast, *cas6* expression was not influenced (Figure 1C). To confirm this effect derived from CreA rather than CreT mutation, we examined the fluorescence from *P_creT_* or P_*cas8*_-controlled *gfp* in WT, Tm and TAdm cells. As expected, fluorescence was not (P_*creT*_) or only slightly (P_*cas8*_) increased by CreT mutation (possibly due to the translation-inhibiting effect of CreT toxin (Li et al., 2021)), while for both promoters, fluorescence greatly elevated (by > 10-fold) in TAdm where CreA was further mutated (Figure 1D). Then we introduced single nucleotide substitutions into the target site of CreA, after which *P_cas8_* and P_*creT*_ both became de-repressed in WT or Tm cells (Figure 1E), reaffirming that they are synchronously downregulated by CreA.

### CreA-guided Cas repression reduce autoimmune risks

The upregulated *cas* genes in TAdm imply that this mutant be more proficient in CRISPR immunity. We introduced into Tm and TAdm cells a synthetic mini-CRISPR with a 34-bp spacer (namely v10) targeting the *Haloarcula hispanica* pleomorphic virus 2 (HHPV-2) virus (Li *et al*., 2014; Gong *et al*., 2019), and then subjected them to HHPV-2 infection. We observed equivalent viral immunity for these two hosts (Figure 2A), suggesting the wild expression level of Cas proteins (in Tm) was sufficient to provide robust immunity. We previously revealed that reducing the size of the spacer or the 3’ handle component of crRNAs would compromise their immunity effects (Gong *et al*., 2019). So, we designed v10-crRNA variants with a truncated spacer or shortened 3’ handle, and found that TAdm was more resistant to HHPV-2 infection than Tm when v10 spacer was truncated to 31 or 32 bp, or when 3’ handle was no more than 10 bp (Figure 2A). Therefore, under some circumstances, TAdm does possess stronger CRISPR immunity.

**Figure 2.**
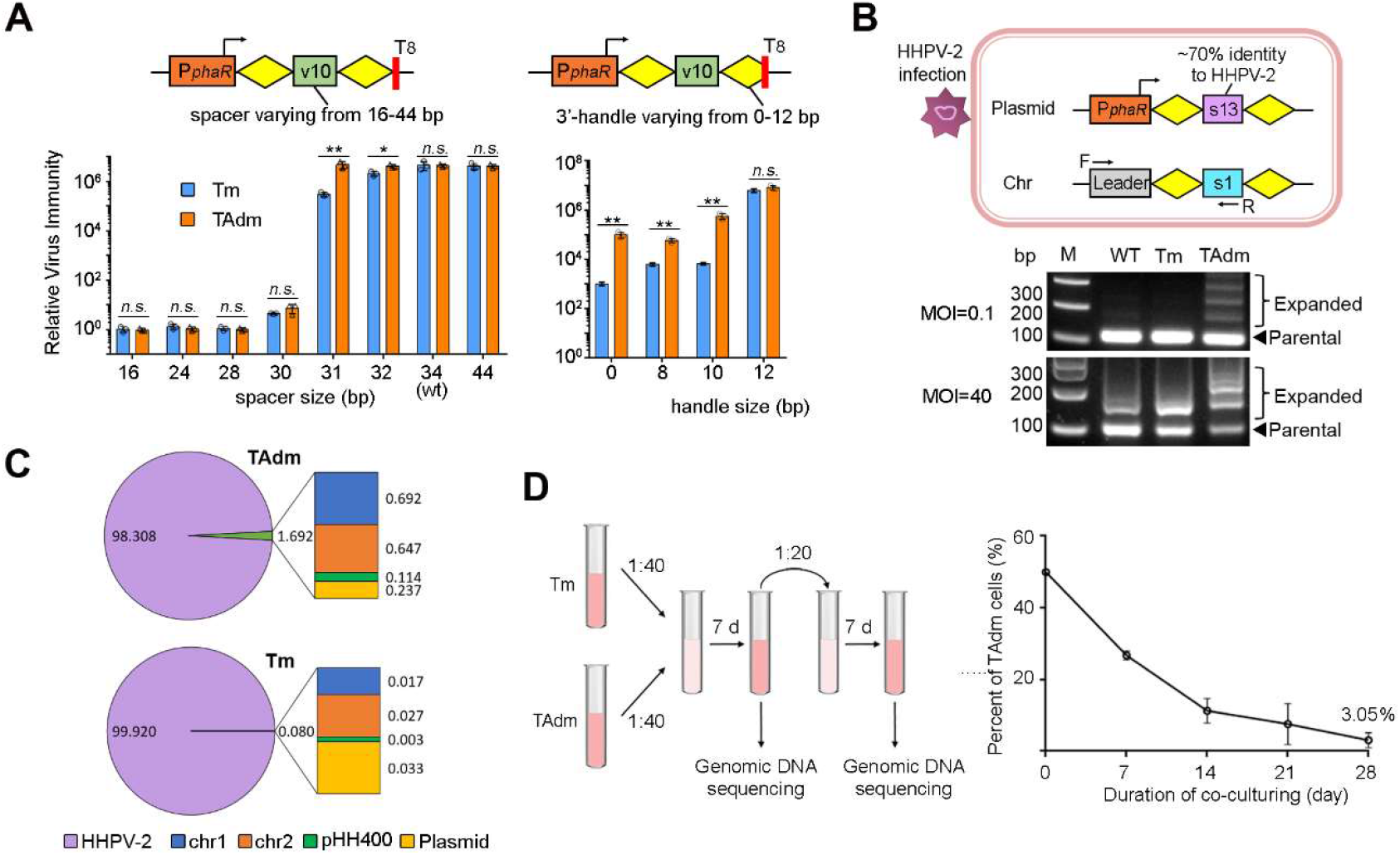
TAdm showed stronger viral immunity with the risk of autoimmunity. (**A**) The virus immunity conferred by different variants of a virus-targeting crRNA (v10) in Tm and TAdm. A constitutive promoter (*P_phaR_*) was used to express the artificial mini-CRISPR, and a string of eight thymines (T_8_) was used as a terminator. Two-tailed Student’s *t* test [*P < 0.05; \*\**P* < 0.01; *n.s*., not significant (P > 0.05)]. (**B**) CRISPR adaptation to HHPV-2 in Tm or TAdm cells over-expressing the s13-crRNA (partially complementary to the viral DNA). Primers specific to the chromosomal CRISPR was used for PCR amplification and the expanded PCR products indicated acquisition of new spacers. MOI, multiplicity of infection. **(C)** The origin ratio (%) of new spacers. *H. hispanica* genome consists of two chromosomes (chr1 and chr2) and one mega-plasmid (pHH400). (**D**) Co-cultivation of Tm and TAdm cells. Cell percent was analyzed by high-throughput DNA sequencing. Error bars, mean±s.d. (n=3). See also Figure S3 and S4.

Because the core *cas* genes involved in adaptation (*cas1,cas2*, and *cas4*) were also upregulated in TAdm, we inferred this mutant be more effective also in acquiring new spacers. The *H. hispanica* CRISPR array has 13 spacers, and the terminal one (s13) shares ~70% sequence identity to HHPV-2 and primes efficient spacer acquisition from this virus (Li *et al*., 2014). Regarding that the CRISPR array might be also upregulated in TAdm (though we did not detect marked increase in crRNA abundance in this strain, see (Figure S3A), we engineered a s13-crRNA-expressing CRISPR to ensure equivalent amounts of priming molecules in WT, Tm and TAdm cells (Figure 2B), which was further verified by Northern blotting (Figure S3B). When infected at a low or high MOI (multiplicity of infection, 0.1 or 40), TAdm, as expected, incorporated new spacers into the chromosomal CRISPR array more frequently than WT and Tm (Figure 2B). However, by Illumina sequencing, we found that TAdm mistakenly acquired self-derived spacers at a frequency of ~1.69%, which occurred much more rarely in Tm (~0.08%) (Figure 2C). We speculated most (if not all) of these self-derived spacers were acquired via naïve adaptation that not relies on a priming step. In accordance, we observed that, after long-term cultivation without virus infection, TAdm cells very inefficiently acquired endogenous DNA as new spacers (Figure S4). Therefore, CRISPR adaptation does tone up in TAdm, but with a risk of spawning self-targeting spacers via the naïve pathway. We predicted the accumulation of self-targeting spacers (and possibly also the high expression of Cas proteins) should compromise the fitness of TAdm cells. Consistently, when we cocultured Tm and TAdm for a long period, the former outcompeted the latter, and after 28 days, the proportion of TAdm cells descended to 3.05% (Figure 2D).

### CreA-repressed promoters can sense crRNA abundance

Presuming that crRNA and CreA RNA compete for Cas proteins to form an immune or regulatory effector, we inferred that the CreA-guided gene repression could respond to alterations in crRNA amounts. Because Cascade-CreA possibly not only suppresses the activity of *P_cas8_*, but also attenuates the potential readthrough transcripts driven by *P_cas6_*, we probed the transcription level of *cas8* relative to *cas6* in *H. hispanica* WT and ΔCRISPR cells. As expected, in ΔCRISPR, the relative RNA abundance of *cas8* declined to ~80% of WT level (Figure S5). A similar decline in *cas8* transcription was observed for a CRISPR mutant encoding only one spacer (Δsp2-13; Figure S5). Then we introduced the *P_cas8_* or P_*creT*_-controlled *gfp* into these cells, and found that, for either promoter, fluorescence declined by 10-20% in ΔCRISPR and Δsp2-13 mutants compared to WT (Figure 3A), which directly illustrated that P_*cas8*_ and P_*creT*_ were more tightly repressed by CreA when CRISPR volume decreased. We further constructed a ΔCRISPR mutant based on Tm (namely Tm-ΔCRISPR), and then supplemented a copy of the original CRISPR using a plasmid. Cells containing a leader-preceded or a *P_phaR_*-driven CRISPR array produced *cas8* transcripts almost twice as much as the cells containing a leader-less array (Figure 3B), illustrating that CreA-guided gene repression was relaxed by enlarging the crRNA pool.

**Figure 3.**
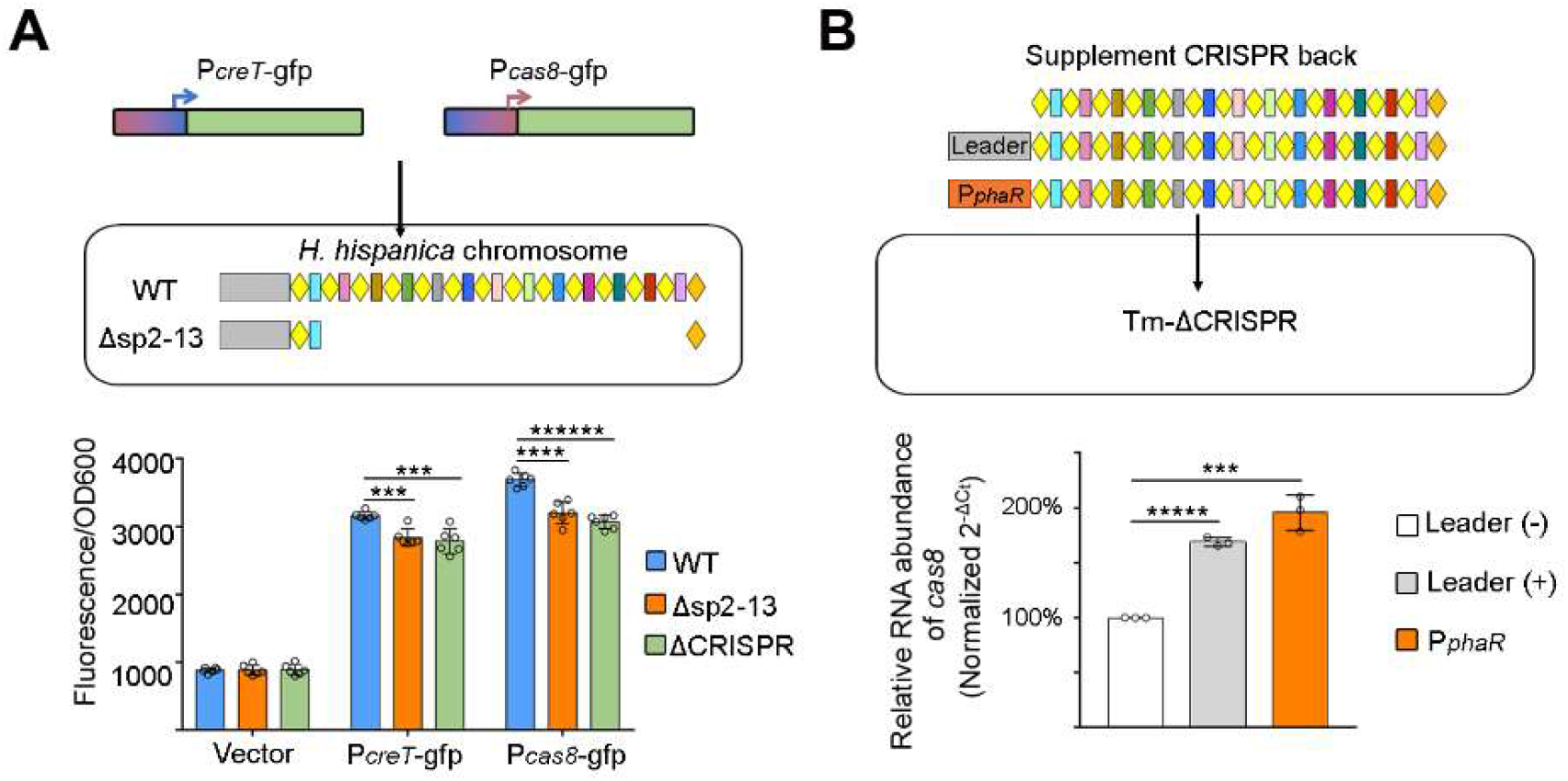
*P_creT_* and *P_cas8_* sense CRISPR volume. (**A**) Activity of *P_creT_* and *P_cas8_* in cells containing 13 (WT), 1 (Δsp2-13) or 0 (ΔCRISPR) CRISPR spacers. (**B**) Relative RNA abundance of *cas8* in cells lacking (leader-) or expressing crRNAs (leader+ or *P_phaR_*). RNA of *cas6* served as the internal control. Tm-ΔCRISPR, a CRISPR-minus mutant constructed based on Tm. Error bars, mean±s.d. (n=3); two-tailed Student’s *t* test [\*\*\**P* < 0.001; \*\*\*\**P* < 0.0001; \*\*\*\*\**P* < 0.00001; \*\*\*\*\*\**P* < 0.000001]. See also Figure S5.

### Type I CRISPR-Cas is widely regulated by degenerated crRNAs

Then we asked whether Cas expression is widely repressed by CreA or other CreA-like degenerated crRNAs. By revisiting the previously discovered *creTA* analogs associating with I-B CRISPR systems (Li *et al*., 2021), we found that, in several cases, the target site of CreA locates within or next to the putative promoter of their *cas* operon (Figure S6). By manually searching the *cas* intergenic sequences of more CRISPR-Cas systems, we also found some creA-like elements (degenerated mini-CRISPRs) for I-D, I-C, I-E, I-F, and I-U subtypes (Figure S7), and in most cases, the target site of CreA closely adjoins the predicted promoter controlling the effector *cas* genes. Notably, a considerable portion of these elements seemingly not cooccur with a toxic gene, like the case of a *Salmonella enterica* I-E CRISPR-Cas locus (illustrated in Figure 4A). For convenience while avoiding confusions, we propose to refer to these seemingly ‘ standalone’ *creA* genes as *creR* (<u>C</u>as-<u>re</u>gulating <u>R</u>NA) before their coupling toxin genes are convincingly identified or predicted. It appears that Cas autoregulation guided by CreA or CreR is a general mechanism of different type I CRISPR systems.

**Figure 4.**
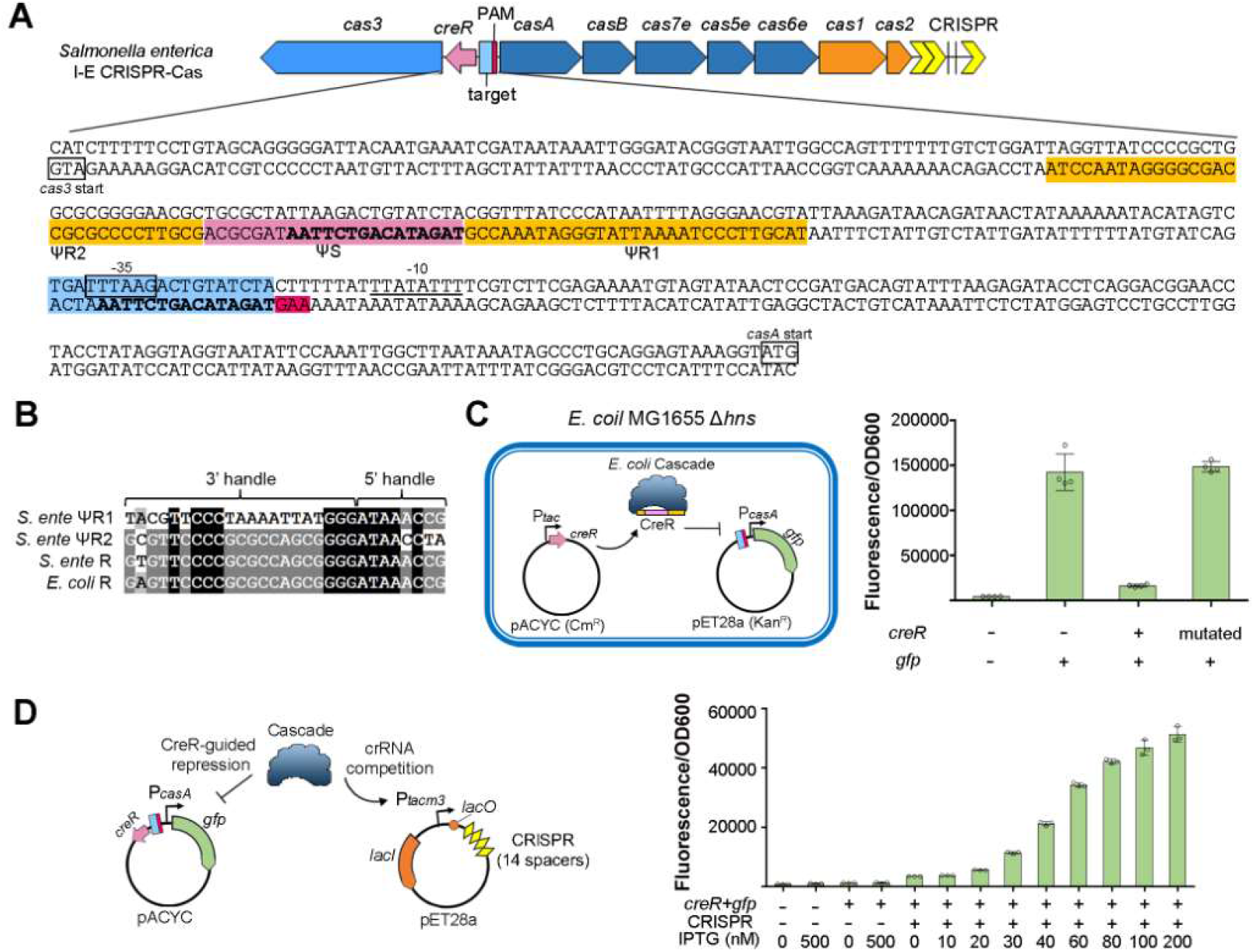
The S. *enterica* I-E CRISPR-Cas employs a degenerated mini-CRISPR (namely *creR*) to regulate Cas expression. (**A**) Schematic depiction of the CRISPR-Cas architecture and the location and sequence of *creR*. Promoter elements (−35 and −10) of *casA* were predicted. Nucleotides in bold indicate the identical ones shared between the ‘spacer’ (ΨS) of CreR and its corresponding protospacer. (**B**) Homology between the CRISPR repeat (R) and the CreR ΨR sequences. The CRISPR repeat of *E. coli* MG1655 is included for comparison. (**C**) S. *enterica* CreR reprogramed *E. coli* Cascade to repress the cognate P*_casA_*. The first 3 nucleotides (TAG) of the CreR ΨS sequence was mutated to disrupt its complementarity to P_*casA*_. Error bars, mean±s.d. (n=4). (**D**) CreR-guided repression of P*_casA_* was relieved by over-expressing the competing crRNAs. The CRISPR array of *E. coli* DH5α was amplified and over-expressed. Error bars, mean±s.d. (n=3). See also Figure S6 and S7.

Because the CRISPR repeat of the *S. enterica* ATCC 51960 system is highly similar (differs by only one nucleotide) to that of the *Escherichia coli* MG1655 CRISPR-Cas system (Figure 4B), we selected to test the Cas repression effect of *S. enterica* CreR in this *E. coli* strain (Figure 4C). Similar to the case of *creA*, ΨR1 of this *creR* gene holds more conservation for the 8 nucleotides that produce a 5’ handle on the mature RNA, while ΨR2 holds more conservation for those corresponding to a 3’ handle (Figure 4B). The ΨS sequence share 15 consecutive nucleotides with its target site (flanked by a 5’-AAG-3’ trinucleotide, i.e., the canonical PAM of I-E systems), which locates ~140 bp upstream of *casA*, the first gene of the *cascade* operon (Figure 4A). Therefore, we amplified a long promoter sequence (206 bp) of *S. enterica casA* (referred as P*_casA_*) to include this target site and put *gfp* under its control. We observed that fluorescence intensity declined by ~8.6-fold when CreR was *in trans* produced (driven by the commonly used *tac* promoter) from another plasmid (Figure 4C). As expected, this repression effect was lost when we further mutated CreR to disrupt its complementarity to P*_casA_*. Therefore, the *S. enterica* CreR repressed P*_casA_* in the help of the *E. coli* Cascade, which indicates the I-E Cascade effector in *S. enterica* employs this degenerated crRNA to achieve its autorepression.

### CreR-regulated Cas expression is positively finetuned by crRNA volume

Utilizing the well-developed genetic tools in *E. coli*, we tested and characterized the correlation between the activity of CreR-repressed *cas* promoter and the volume of crRNA molecules in the cell (Figure 4D). We engineered the *S. enterica creR* gene and the P_*casA*_-controlled *gfp* into one plasmid, and then expressed a 14-spacer CRISPR array (cloned from the *E. coli* DH5α strain) from another plasmid, using an IPTG-inducible promoter (containing the *lacO* operator). Note that a less active mutant of *tac* promoter (*tacm3*) (Zhang *et al*., 2016) was used to control CRISPR to avoid high level of basal expression. Exponential MG1655 cells containing both plasmids were induced by adding different doses of IPTG. Notably, fluorescence intensity was observed to be increased by a factor of 1.6, 3.3, 6.1, 9.8, 12.0, 13.3, and 14.6, when crRNA production was induced by 20, 30, 40, 60, 80, 100, and 200 nM IPTG, respectively (Figure 4D), which illustrated their positive correlation. It was indicated that the cellular concentration of crRNA molecules, which compete with CreR for Cas proteins, could fine tune the CreR-mediated Cas autorepression to meet their own need of protein partners.

### CreR-guided autorepression of the V-A effector Cas12a

To further extend the generality of the Cas autorepression circuit, we sought to manually search for *creR* (or *creA*) elements in class 2 CRISPR-Cas systems. Intriguingly, from the upstream sequence of *cas* operons, we did obtain a dozen of putative *creR* genes associating with V-A systems, and in the most cases, their predicted target sites locate within or next to the putative promoter of the corresponding *cas12a* (Figure S8), which encodes the V-A single-protein effector (Zetsche *et al*., 2015). For experimental validation in *E. coli* cells (MG1655), we selected the *creR* gene associating with the well-studied *Moraxella bovoculi cas12a* (Mb*_cas12a_*) (illustrated in Figure 5A). This *creR* gene carries a sequence (ΨR1) that is considerably similar to its cognate CRISPR repeat and another extensively degenerated ‘repeat’ (ΨR2) that share very little nucleotide identity with them (Figure 5B). In addition, the RNA of ΨR1 and CRISPR repeat can form a similar stem-loop structure (Figure 5C) and share 20 consecutive nucleotides that give rise to an identical 5’ handle on their mature RNAs (Figure 5B), which strongly suggests a crRNA-like architecture of mature CreR. The spacer portion of *creR* partially complements to a target site that is flanked by a 5’-TTTA-3’ motif (a typical PAM of V-A subtype) and locates very adjacent to the predicted −35 element of the promoter of *cas12a* (P*_cas12a_*) (Figure 5A). We also performed primer extension assay to determine the transcription start site of *P_cas12a_* (Figure S9), which confirmed the prediction of −35 element. Then we synthesized a 330-bp DNA construct including *creR* (and its putative promoter) and *P_cas12a_*, and put *gfp* under its control (Figure 5D). In MG1655 cells, the plasmid carrying this DNA construct produced green fluorescence, which could be markedly suppressed (by ~33-fold) when we introduced another plasmid to *in trans* provide MbCas12a (Figure 5D). This repression effect disappeared when we mutated the spacer portion (ΨS) of *creR* or its target site within *P_cas12a_*, and notably, persisted when *creR* and P_*cas12a*_ were complementarily mutated at the same time. Therefore, based on their partial complementarity, the *M. bovoculi* CreR repressed P_*cas12a*_ in the help of Cas12a in *E. coli*, which indicate a CreR-guided autorepression circuit of this V-A effector in its native host *M. bovoculi*.

**Figure 5.**
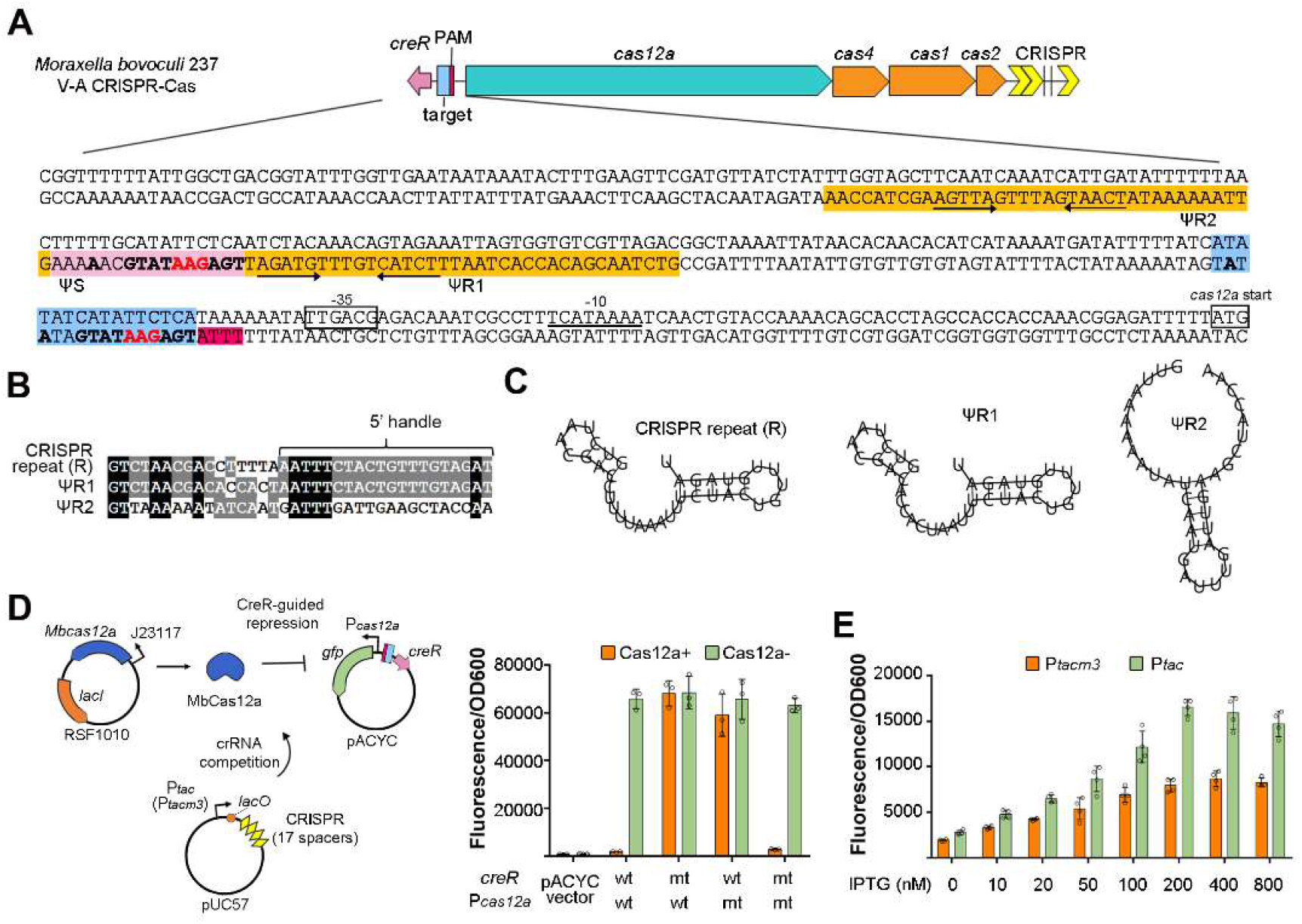
The *M. bovoculi* V-A CRISPR-Cas also employs CreR to regulate Cas expression. (**A**) Schematic depiction of the CRISPR-Cas architecture and the location and sequence of *creR*.Promoter elements (−10 and −35) of *cas12a* were predicted. Nucleotides in bold indicate the identical ones shared between the ‘spacer’ (ΨS) of CreR and its target site. The convergent black arrows indicate inverted repeats within ΨR sequences. (**B**) Homology between the CRISPR repeat (R) and the CreR ΨR sequences. Nucleotides corresponding to the 5’ handle remaining on mature RNA are indicated. (**C**) The stem-loop structure predicted for CRISPR repeat and ΨR sequences. **(D)** Validation of the CreR-guided repression circuit of MbCas12a in *E. coli*. Nucleotides 4-6 in the ΨS sequence of *creR* (indicated in red in panel **A**) and their corresponding nucleotides in the target in P*_cas12a_* were separately or simultaneously mutated (mt). wt, wild-type. MbCas12a was expressed from a plasmid containing a broad host range replication origin (RSF1010) and controlled by J23117 promoter. For the assay in panel **E**, pUC57 was further utilized to express crRNAs. The *M. bovoculi* CRISPR array containing 17 spacers was synthesized. (**E**) The effects of varying crRNA doses on CreR-guided gene repression. Error bars, mean±s.d. (n=3). See also Figure S8 and S9.

Next, we introduced a third plasmid expressing crRNAs to assess their effects on *M. bovoculi* CreR-guide repression (Figure 5D). The CRISPR array of *M. bovoculi* 237, which contains 17 spacers, was synthesized and placed under the control of an IPTG-inducible promoter. In exponential MG1655 cells containing all the three plasmids illustrated in Figure 5D, the fluorescence increased by a factor of 1.7, 2.2, 2.8, 3.6, 4.1, and 4.5 when crRNA production was under control of *P_tacm3_* and induced by 10, 20, 50, 100, 200, and 400 nM IPTG (Figure 5E), respectively, which illustrated a positive correlation between them. When more IPTG (800 nM) was added, fluorescence did not further increase, suggesting that the cellular concentration of crRNA or its relieving effect on CreR repression reached saturation. We propose that it was the crRNA concentration (or its promoter activity) rather than the relieving effect reached saturation, because when we changed P_*tacm3*_ to *P_tac_*, the fluorescence further increased by nearly 1.5 to 2.0-fold at the same induction intensity (i.e., IPTG concentration) (Figure 5E). Therefore, we conclude that the CreR-guided autorepression of Cas12a in *M. bovoculi* can sense and respond to variations in the cellular concentration of crRNA molecules.

### Acr proteins can relieve or subvert CreR-guided Cas autorepression

To defeat against the diverse CRISPR-Cas systems, phages have evolved at least equally diverse small anti-CRISPR (Acr) proteins (Borges *et al*., 2017; Pawluk *et al*.,2018). We further investigated how the Cas autorepression respond to the action of Acr proteins. For the I-E Cascade of MG1655, only one Acr protein, i.e., AcrIE8, has been reported (Hicks *et al*., 2019). So, we synthesized the gene encoding this protein and expressed it using an IPTG-inducible *tacm3* promoter. In exponential MG1655 cells containing the plasmid expressing AcrIE8 and another plasmid carrying the *S. enterica creR* gene and a *gfp* gene controlled by its cognate P*_casA_*, we measured the fluorescence intensity after different doses of IPTG was added (Figure 6A). Notably, fluorescence was increased by a factor of 11.3, 25.5, 28.9, 29.3, and 31.9 when AcrIE8 expression was induced by 10, 20, 30, 40, and 60 nM IPTG, respectively, and was not further increased when more IPTG was used. It appears that a low expression level of AcrIE8 could effectively relieve or even subvert the repression effect on P*_casA_*, which indicates the Cas autorepression circuit can actively respond to the action of Acr proteins to elicit mass production of new Cas weapons.

**Figure 6.**
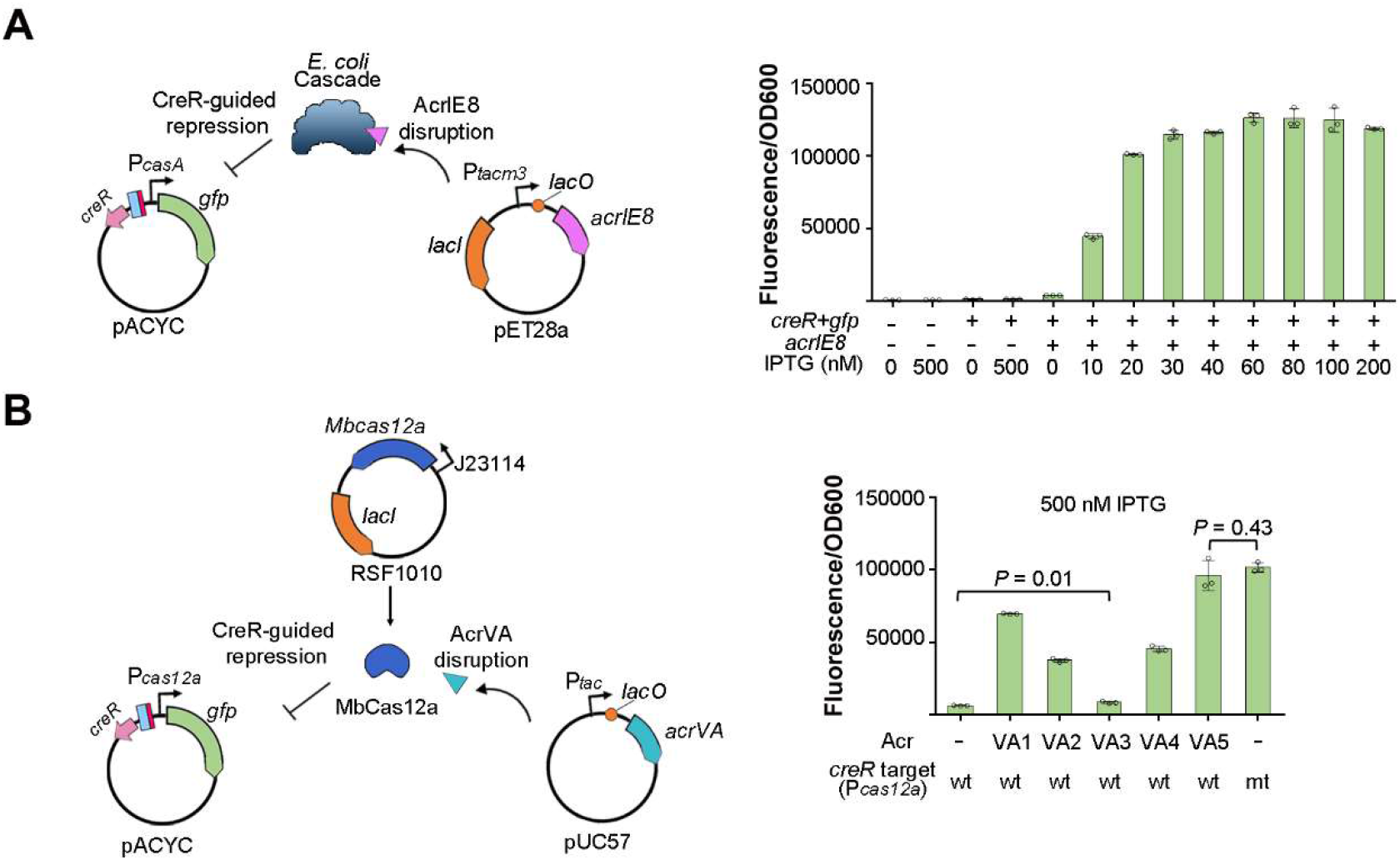
CreR-guided Cas autorepression can be subverted by diverse Acr proteins. (**A**) Disrupting the autorepression circuit of I-E Cascade by expressing the AcrIE8 protein (reported to inactivate MG1655 Cascade). **(B)** Disruption the autorepression circuit of MbCas12a by expressing diverse AcrVA proteins. Error bars, mean±s.d. (n=3). P values were calculated from two-tailed Student’s *t* test. See also Figure S7 and S8.

For the V-A effector Cas12a, at least five Acr proteins (named AcrVA1-5) have been reported (Watters *et al*., 2018; Marino *et al*., 2018), which allowed us to interrogate the effects of different Acr proteins on a same Cas autoregulation circuit. Similar to the crRNA induction assay (depicted in Figure 5D), we employed three plasmids: one plasmid to carry *M. bovoculi creR* and P_*cas12a*_-controlled *gfp*, the second plasmid to encode the MbCas12a effector, and the third one to produce AcrVA proteins under the control of an inducible promoter (Figure 6B). In exponential MG1655 cells containing all these three plasmids, we found that the CreR-repressed fluorescence was markedly but very differently relieved by AcrVA1-5 proteins (Figure 6B). When 500 nM IPTG was used to induce Acr expression, AcrVA3 only increased the fluorescence intensity by 1.3-fold, while AcrVA1, AcrVA2, and AcrVA4 increased by a factor of 10.9, 5.9, and 7.2, respectively, with AcrVA5 showing the strongest increasing effect (by 15.1-fold), which appeared to be equivalent to the effect of disrupting the complementarity between CreR and P_*cas12a*_ (Figure 6B). Therefore, the autoregulation circuit of MbCas12a respond quite differently to various AcrVA proteins, suggesting their different detailed anti-CRISPR mechanisms. In fact, it was reported that these AcrVA proteins showed varying inhibiting effects on Cas12a proteins from different *M. bovoculi* strains.

## DISCUSSION

There are increasing evidences supporting that CRISPR-Cas effectors have a secondary physiological role in regulating host genes, in addition to its canonical immune function. The type II effector Cas9 was demonstrated to be reprogrammed by a small CRISPR-associated RNA (scaRNA) to regulate a virulence-related gene that encodes a lipoprotein (Ratner *et al*., 2019). Our group has recently performed systemic investigations into the addiction module of CRISPR-Cas, i.e., CreTA, wherein the antitoxin gene *creA* actually evolved from degenerated mini-CRISPRs and reprograms the type I effector Cascade to repress the expression of a toxic RNA (CreT) (Li *et al*., 2021; Cheng *et al*., 2021; Cheng *et al*., 2022). In this study, we further showed that CreA and its CreR analogs (not cooccurring with a toxin gene) widely distribute in type I and V systems (assigned to class 1 and class 2, respectively (Makarova *et al*., 2020)), and provided experimental evidences that these regulatory degenerated CRISPRs commonly mediate the autorepression of both multi-subunit (I-B and I-E) and single-protein (V-A) Cas effectors. Notably, this systematic discovery complements the recently unraveled autorepression mechanism of Cas9 where a long isoform of tracrRNA (trans-activating CRISPR RNA) seemingly plays the role of CreR (Workman *et al*., 2021). Therefore, it seems to be a general paradigm for both class 1 and class 2 CRISPR-Cas systems that the Cas effectors are repurposed by noncanonical RNA guides for their autoregulation.

We propose the Cas autorepression circuit exquisitely balances the immune benefits and the fitness costs of CRISPR-Cas. To provide effective immunity, a sufficient production level of Cas proteins is theoretically required to satisfy the need of crRNA guides, which is ever changing during the conflicts between bacteria and phages. Moreover, when infected by phages encoding small Acr proteins that inactivate the Cas effector, there will be an urgent need of mass production of new Cas proteins to quickly reboot CRISPR immunity. However, a constant high expression level of the multi-subunit (class 1) or high molecular weight (class 2) Cas effector will cause considerable energy costs and possibly result in deleterious autoimmune events (as observed in Figure 2C where Cas autorepression was eliminated), which will certainly lower down the fitness of host cell (Figure 2D). The Cas autorepression circuit, which involves Cas proteins *per se* and a non-canonical guide (CreR or CreA) that competes with crRNAs, makes Cas expression responsive to Acr elements that attack Cas proteins and also to alterations in the cellular concentration of crRNAs (Figure 7). Note that, the repressed *cas* promoters need be highly effective to support mass production of Cas proteins when required. Consistently, in the absence of CreA, the *cas* promoter (P_*cas8*_) in *H. hispanica* turned out to be the most effective haloarchaeal promoter we have ever tested (Figure S2). From the view of arms race, Cas autorepression may represent a new anti-anti-CRISPR strategy that acts on transcriptional level.

**Figure 7.**
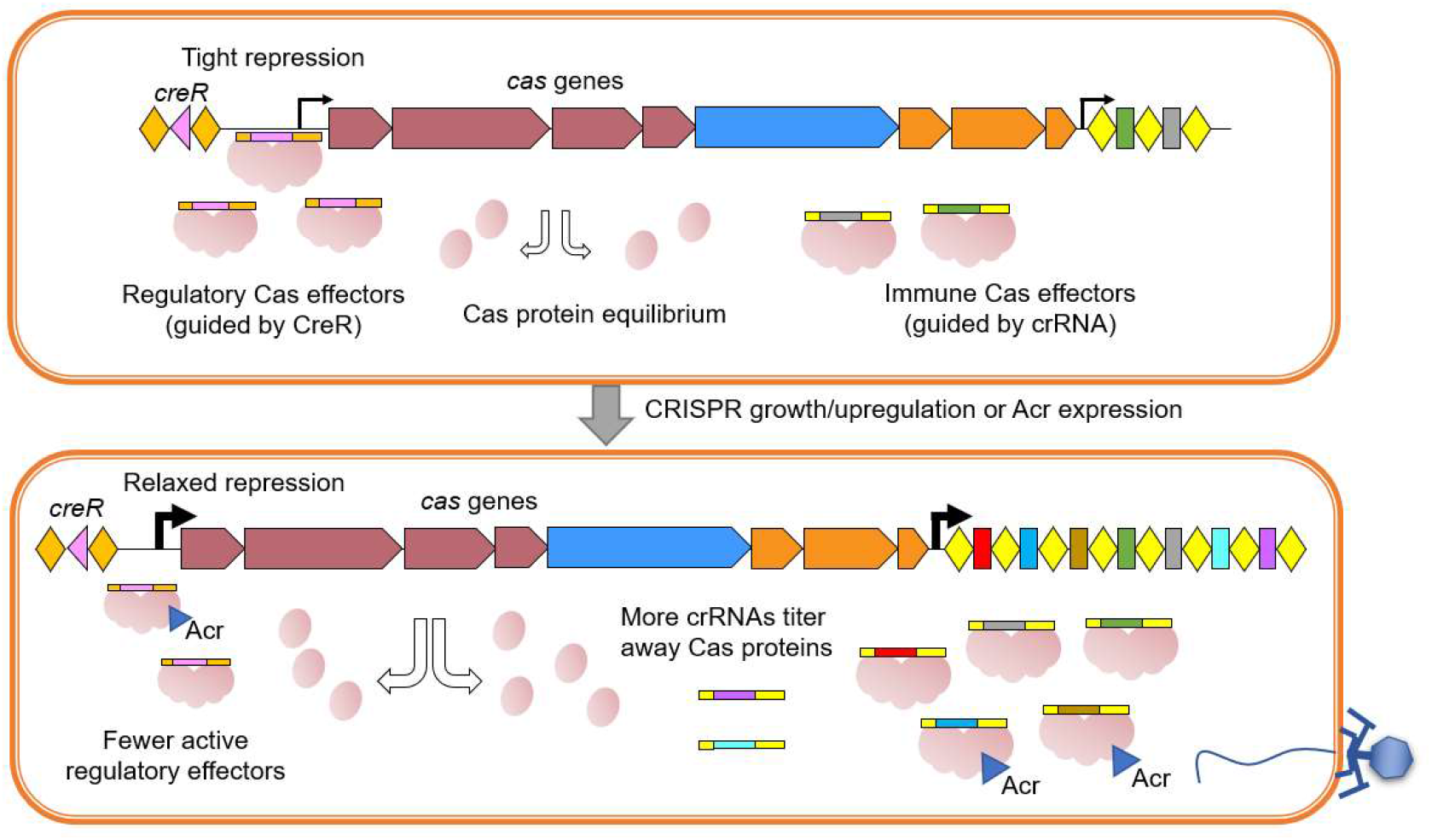
The model of CreR-guided Cas autorepression. In cells with a low level of crRNA production, CreA guides Cas effectors to tightly repress the promoter of *cas* genes. When the cellular concentration of crRNA elevates, possibly due to the growth or upregulation of CRISPR arrays, Cas proteins are titrated away from the regulatory circuit, and Cas repression gets relaxed to replenish the pool of Cas proteins until a new equilibrium is achieved. When the Acr proteins from an infecting phage inactivate Cas proteins, the Cas autorepression circuit will be completely subverted, leading to mass production of new immune weapons. See also Figure S6, S7 and S8.

During preparation of this manuscript, Shmakov *et al*. reported a systematic search of CRISPR repeat-like RNA regulatory elements (Shmakov *et al*., 2023), which may largely enrich the collection of CreR or CreTA elements. Their study also showed that the expression level of a I-F Cascade was regulated by a predicted CRISPR repeat-like RNA (corresponding to the CreR in this study), which supports and reinforces the generality of Cas autorepression.

In summary, our data provide substantial experimental evidences for the Cas autoregulation circuit, which intriguingly, is directed by degenerated crRNAs (CreR or CreA), and remarkably, is responsive to the changing need of typical defensive crRNAs and susceptible to the action of Acr proteins. We propose this regulation circuit can exquisitely balance the immune competence and the fitness cost of CRISPR-Cas, and, in combination with the diverse CRISPR-regulated toxins (e.g., CreT), likely have promoted the wide distribution and stable persistence of CRISPR-Cas in prokaryotes.

## MATERIALS AND METHODS

### Strains and growth conditions

*H. hispanica* DF60 (uracil auxotroph mutant of *H. hispanica* ATCC 33960) (Liu *et al*., 2011) or its derivatives were cultivated at 37°C in nutrient-rich AS-168 medium (per liter, 200 g NaCl, 20 g MgSO_4_·7H_2_O, 2 g KCl, 3 g trisodium citrate, 1 g sodium glutamate, 50 mg FeSO_4_·7H_2_O, 0.36 mg MnCl2·4H_2_O, 5 g Bacto Casamino Acids, and 5 g yeast extract, pH 7.2) supplemented with uracil at a final concentration of 50 mg/liter. For the strains carrying the expression plasmid pWL502 or its derivatives, the AS-168 medium with yeast extract subtracted was used. All strains were cultivated either on solid agar plates (1.2% agar) or in liquid cultures.

*E. coli* DH5α was used for plasmid construction. *E. coli* MG1655 and its mutants were used for CreR-guided repression assays of I-E and V-A type CRISPR-Cas. All bacterial strains were grown at 37°C in Luria-Bertani (LB) medium (per liter, 10 g tryptone, 5 g yeast extract, 10 g NaCl), with agar (12 g/L) for solid plates and with 200 rpm shaking for liquid cultures. When needed, antibiotics were added to the following concentrations: apramycin sulfate (50 μg/mL), ampicillin (100 μg/mL), kanamycin (50 μg/mL), or chloramphenicol (25 μg/mL).

### Purification and concentration of the virus virions

The top agar of a single plaque containing the HHPV-2 virions was inoculated into an early exponential culture of *H. hispanica* for enrichment. After 5-day cultivation (37°C, 200 rpm), the culture was collected and the cells were removed by centrifugation (9,000 rpm, 4°C, 15 min). The supernatant was subjected to the VIVAFLOW 50 system (Sartorius, 50,000 MWCO) for pre-purification and subsequently to the 0.22 μm filters.

### Plasmid construction

Plasmids, oligonucleotides and synthetic gene used in this study are listed in Table S1. The double-stranded DNA fragments were amplified using the high-fidelity KOD-Plus DNA polymerase (TOYOBO, Osaka, Japan), then digested by restriction enzymes (New England Biolabs, MA, USA) and ligated into the predigested vector using the T4 DNA ligase (New England Biolabs, MA, USA), or directly assembled into predigested plasmids through Gibson assembly strategy using Trelief^®^ Seamless Cloning Kit (Tsingke Biological Technology, Beijing, China). Mutant construction was performed using the overlap extension PCR strategy. The engineered plasmids were validated by DNA sequencing.

### Transformation

For plasmids introduced into haloarchaeal cells. Transformation of the haloarchaeal strains was performed using the polyethylene glycol-mediated method according to the online Halohandbook (https://haloarchaea.com/wp-content/uploads/2018/10/Halohandbook_2009_v7.3mds.pdf). The yeast extract-subtracted AS-168 plates were used to screen the transformants. The average and standard deviation of transformation efficiency (colony forming unit per μg plasmid DNA, CFU/μg) were calculated based on log-transformed data.

Regarding the introduction of plasmids into *E. coli* MG1655, the electro-transformation method was utilized. The bacteria were cultured in 10 mL LB broth at 37°C overnight with 200 rpm shaking. Subsequently, the cultures were sub-inoculated into 200 mL fresh medium (1:100 dilution) and grown to an OD_600_ of 0.6. The cells were then collected via centrifugation at 4 °C, washed thrice using cold double-distilled water, and finally resuspended into 2 mL of ice-cold water containing 10% glycerol. A total of 50 μL of cells was mixed with plasmids and then subject to electroporation (Bio-Rad, USA) at 2.5 kV. The shocked cells were recovered with 1 mL LB at 37°C for 1 h, and then plated on LB plates containing appropriate antibiotics. In cases where multiple plasmids were required, the method could be repeated via multiple electrical transfers.

### Mutant construction and gene knockout

Plasmids and oligonucleotides used for mutant construction in this study are listed in Table S1. Mutant construction or gene knockout in haloarchaeal cells was performed as previously described (Liu *et al*., 2011). For example, to construct the TAdm mutant, the Shine-Dalgarno (SD) motif of *creT* and the first two nucleotides of the CreA seed sequence in *creTA* (NC_015943.1: 145387-145697) were mutated by PCR amplification, its upstream ~500 bps and downstream ~500 bps were separately amplified, then all the three fragments were connected by overlap extension PCR using the corresponding primer pairs. The linked fragment was digested and inserted into the suicide plasmid pHAR (Liu *et al*., 2011). The constructed plasmids were validated by DNA sequencing and then transformed into *H. hispanica* ΔTA cells. After screening of single and then double cross-over mutants, the TAdm mutant cells were validated by colony PCR and subsequent Sanger sequencing.

Mutants or gene knockouts in *E. coli* MG1655 were generated using the λ red and flp/FRT system, as described in previous studies (Doublet *et al*., 2008; Luo *et al*., 2015). For example, to create the mutant *E. coli* MG1655(Δhns, *Δcas3* & *P_J23119_*), a fragment containing the upstream 40bp, followed by FRT-Kan^R^-FRT-P_*J23119*_ sequence, and downstream 40bp was electroporated into *E. coli* MG1655, which was previously transformed with pKD46 and induced with 1% L-arabinose. The kanamycin-resistant colonies were screened and verified using PCR, and then the kanamycin resistance gene was removed with pCP20 and a temperature shift to 42°C. The resulting mutant cells were further confirmed by colony PCR and Sanger sequencing.

### Fluorescence measurement

To test the promoter activity for type I-B CRISPR-Cas system in haloarchaeal strains, we firstly constructed the plasmid containing the GFP (Reuter *et al*., 2004) based on the pWL502 plasmid. By overlap extension PCR, promoter DNA and its mutated sequences were linked to the gene of a soluble-modified red-shifted version of GFP. The expression vector pWL502 with this hybrid DNA was then introduced into the *H. hispanica* cells by using the above transformation methods.

To test the promoter activity in bacterial stains for I-E and V-A type CRISPR-Cas system, the native chromosome I-E CRISPR-Cas system of *E. coli* MG1655(Δhns) was used in Figure 4C, the *cascade* over-expression mutant strain *E. coli* MG1655(Δhns, *Δcas3* & *P_J23119_*) was used in Figure 4D & 6A, and the *E. coli* MG1655 with *P_J23117_* promoter-controlled Mbcas12a plasmid was used for V-A type CRISPR-Cas system in Figure 5D, 5E & 6B. The 330-bp DNA upstream of *M. bovoculi cas12a* or 206 bp upstream of *S. enterica casA* (*creR* contained sequences), and the GFP-mut3 gene were amplified and assembled into pACYC vector. Subsequently, the GFP reporter plasmid and its mutated plasmids were electroporated into the above *E. coli* MG1655 or its mutants.

To test the promoter activity under the influence of IPTG-induced CRISPR or Acr, 30 μL of bacteria in the late exponential phase were transferred to a 3mL fresh LB culture containing suitable antibiotics. Additionally, 0.5mM or varying amounts of IPTG were added as needed. These cultures were cultured for 5 hours, after which their OD_600_ and fluorescence levels were determined simultaneously.

For each sample, at least three individual colonies were randomly selected and cultured with appropriate antibiotics or yeast extract-subtracted AS-168 medium to the logarithmic growth phase, after which their OD_600_ and fluorescence were simultaneously determined using the Synergy H4 Hybrid multimode microplate reader (BioTeck, VT, USA). The OD_600_ and fluorescence of the cultures were simultaneously determined. The fluorescence/OD_600_ ratio was calculated for each of the triplicates, and their average and standard deviation were calculated. Two-tailed Student’s t-test was performed.

### RNA extraction

To extract *H. hispanica* RNA for RT-qPCR and Northern bolt analysis, Transformant colonies were randomly picked and inoculated into 10 mL of As-168 or yeast extract-subtracted AS-168 medium. After sub-inoculation and 2-day cultivation, the total RNA was extracted from exponential cells using the TRIzol reagent (Invitrogen, MA, USA) according to the standard guidelines. RNA concentration was determined using NanoDrop One spectrophotometer (Thermo Fisher Scientific, MA, USA). To extract *E. coli* for primer extension analysis, 30 μL of cultures in the late exponential phase were transferred to a 3mL fresh LB culture containing suitable antibiotics. These cultures were cultured for 5 hours. After the culture had been harvested through centrifugation, TRIzol was added to extract RNA following the procedure described previously.

### qPCR

*H. hispanica* cells were pre-cultured to the early stationary phase and then sub-inoculated into fresh medium and cultured till OD_600_ reached to about 0.8. Cells were collected by centrifugation at 4°C. Total RNA was extracted using the TRIzol reagent (Invitrogen, MA, USA) following the manufacturer’s instructions. A total of 20 μg of RNA was treated with DNase I (Thermo Fisher Scientific, MA, USA) according to the manufacturer’s instructions, and then purified using the phenol: chloroform method. RNA was reverse transcribed into complementary DNA using M-MLV Reverse Transcriptase (Promega, MA, USA). qPCR assay was prepared using KAPA SYBR^®^ FAST qPCR Kit (Kapa Biosystems, MA, USA) and performed on an Applied Biosystems ViiA™ 7 Real-Time PCR System according to the manufacturer’s instruction. The primer sequences used for qPCR are listed in Table S1. For each experimental setting, three biological samples from individual colonies were included, and each sample was examined in triplicate.

### Northern blot analysis

Haloarchaeal cells from 3-5 mL of the early exponential culture were collected by centrifugation. Total RNA was purified using TRIzol (Invitrogen, USA) following the manufacturer’s instructions. A total of 3 μg of RNA was denatured at 65°C for 10 min with equal volume of RNA loading dye (New England Biolabs, MA, USA). RNA samples, a biotin-labeled 64-nt single-stranded DNA and the Century-Plus RNA ladder (Thermo Fisher Scientific, MA, USA) were loaded to an 8% polyacrylamide gel and electrophoresed in 1× TBE buffer. Then, the separated RNAs were transferred onto a nylon membrane (Pall, NY, USA) followed by cross-linking with Ultraviolet (UV) light. The target RNA was hybridized with corresponding biotin-labeled ssDNA probes. The signal was detected using the Chemiluminescent Nucleic Acid Detection Module Kit (Thermo Fisher Scientific, MA, USA) according to the manufacturer’s instructions. The membrane was imaged using the Tanon 5200 Multi chemiluminescent imaging system (Tanon Science & Technology, Shanghai, China).

### Primer extension analysis

The 5’-FAM (6-carboxyfluorescein)-labeled *gfp-specific* primer (Table S1) was ordered from Sangon Biotech (Shanghai) Co., Ltd. 30 μg of the total RNA was firstly digested with RQ1 DNase (Promega, WI, USA), and then reverse transcribed into complementary DNA (cDNA) using 30 enzyme units(U) of the MMLV-RT (Promega, WI, USA) and 1μM of the labeled primer. The extension products were analyzed using the ABI3730xl DNA Analyzer (Thermo Fisher Scientific, MA, USA), and the results were viewed using Peak Scanner Software v1.0.

### Coculturing assay

Tm and TAdm cells were inoculated into 3 ml of fresh AS-168 medium (containing uracil) in triplicate and cultured till OD_600_ 1.0. According to their actual OD values, Tm and TAdm cultures were mixed to nearly equal cell concentrations. 150 μL of the mixture was inoculated into 3 ml of fresh medium and grown for 7 days until stationary phase. Then, Sub-inoculation was performed at a ratio of 1:20 (i.e., inoculating 150 μL of culture into 3 mL of AS-168 medium) every 7 days. Before each sub-inoculation, cells were collected by centrifugation and stored at −20°C. The coculturing persisted for 28 days. To detect the ratio of the total cells of Tm to TAdm, genomic DNA was extracted from the stored cells using the Phenol: Chloroform: Isoamyl alcohol (25:24:1, pH=8.0) method.

A total amount of 0.2 μg DNA per sample was used as input material for the DNA library preparations. Sequencing library was generated using NEB Next^®^ Ultra TM DNA Library Prep Kit for Illumina (NEB, USA, Catalog#: E7370L) following manufacturer’s recommendations and index codes were added to each sample. Briefly, genomic DNA sample was fragmented by sonication to a size of 350 bp. Then DNA fragments were endpolished, A-tailed, and ligated with the full-length adapter for Illumina sequencing, followed by further PCR amplification. After PCR products were purified by AMPure XP system (Beverly, USA). Subsequently, library quality was assessed on the Agilent 5400 system (Agilent, USA) and quantified by QPCR (1.5 nM). The qualified libraries were pooled and sequenced on Illumina platforms with PE150 strategy in Novogene Bioinformatics Technology Co., Ltd (Beijing, China), according to effective library concentration and data amount required.

### Virus interference assay

Individual transformed colonies were randomly selected to inoculate yeast extract-subtracted AS-168 medium. After sub-inoculation and another 2-day culturing, 200 μL of the culture were mixed with 100 μL of 10-fold serial dilutions of the HHPV-2 virus and incubated for 30 min at room temperature. The mix was then mixed with 3 mL of molten 0.7% agar yeast extract-subtracted AS-168 medium at 55°C and immediately poured onto the plates (containing 1.2% agar). Once dry, the plates were incubated for 3 days at 37°C to allow plaque formation. The plaque-forming units (PFU) were counted, and the ratio of the PFU formed on the empty plasmid-carrying strain divided by the PFU formed on the crRNA-expressing strain was used to represent the relative virus immunity (RVI). The average and the standard deviation were calculated based on three replicates.

### Spacer acquisition assay

Individual transformed (by a plasmid over-expressing the crRNA of spacer13) colonies were randomly selected and inoculated into yeast extract-subtracted AS-168 medium. Then, the early-exponential cultures were separately mixed with the HHPV-2 virus at the different MOIs (0.1 and 40). 1 mL of the mixture was inoculated into 10 mL of fresh medium and incubated at 37°C. Spacer acquisition was detected by PCR with specific primers after 1 day.

To monitor the naïve adaptation, individual WT, Tm and TAdm colonies transformed by the empty pWL502 vector were randomly picked and streaked on a fresh plate for further cultivation (at 37°C). Spacer acquisition was examined by colony PCR every week until significant acquisition bands were observed. Three colonies were tested for each strain, and only representative gel images are shown.

### Spacer sequencing analyses

The agarose gel was used to select PCR bands corresponding to the “expanded” a-CRISPR, which were then purified using the efficient AxyPrep™ DNA Gel Extraction Kit from Corning, NY, USA. These purified DNA samples underwent HiSeq2500 sequencing conducted by Biomarker, Beijing, China. The reads containing two or three repeats were selected after assembly of the pair-end data and filtration of low-quality data. For those reads containing two repeats, the intervening sequence was identified as the initially acquired spacer (s-1). For reads with three repeats, the leader-distal new spacer was initially acquired, with the leader-proximal spacer acquired secondly (s-2). Each spacer’s protospacer sequence was preliminarily identified against the HHPV-2 or H. hispanica genome using the BLASTN program, with manual calibration in case of mismatches. The PAM of each protospacer was considered to be the 3 bp 5’-upstream of the sequence. Perl scripts were used to analyze the protospacer sequences and their distribution across the HHPV-2 and DF60 genome (Li *et al*., 2017).

### Bioinformatic analysis

RNA secondary structure was predicted using the RNAfold webserver. Promoter elements were predicted using the BPROM program (Softberry tool).

### Data analysis and image visualization

Microsoft Excel was used to analyze the data, and GraphPad Prism was used to generate the plots. The graphs were then modified in Adobe photoshop to construct the final figures.

## QUANTIFICATION AND STATISTICAL ANALYSES

The number of replicates is specified in the associated figure legends. Each replicate represents a biological replicate of the specified experiment. Two-tailed *t* test was performed for statistical analyses. P-values above 0.05 were considered non-significant. Statistical comparisons for the transformation assays relied on log values, which assumes the samples are normally distributed on a log scale.

## AUTHOR CONTRIBUTIONS

M. L., R. W., and C. L. designed experiments. C. L., R. W., and J. L. constructed mutant strains with the assistance from F. C. and X. S.; C. L. performed the Northern blotting assay, spacer acquisition assay and competition assay with the assistance from S. H. and H. Z.; C. L. and R. W. performed the fluorescence analysis with the assistance from J. L., L. W., J. Y. and Y. Z.; C. L. and F. C. carried out qPCR and transformation assays with the assistance from Q. X. and A. W.; F. C. performed virus interference assay with the assistance from C. L.; M. L., X. S. and H. Y. performed the bioinformatic analyses; M. L. and H. X. analyzed the data and supervised the project. M. L. wrote the manuscript.

## Data and materials availability

All data are available in the main text or the 40 supplementary materials. Reagents are available upon request from M. L.

## SUPPLEMENTARY MATERIALS

Materials and Methods

Figures S1 to S9

Tables S1 to S2

## SUPPLEMENTARY FIGURES

**Figure S1.**
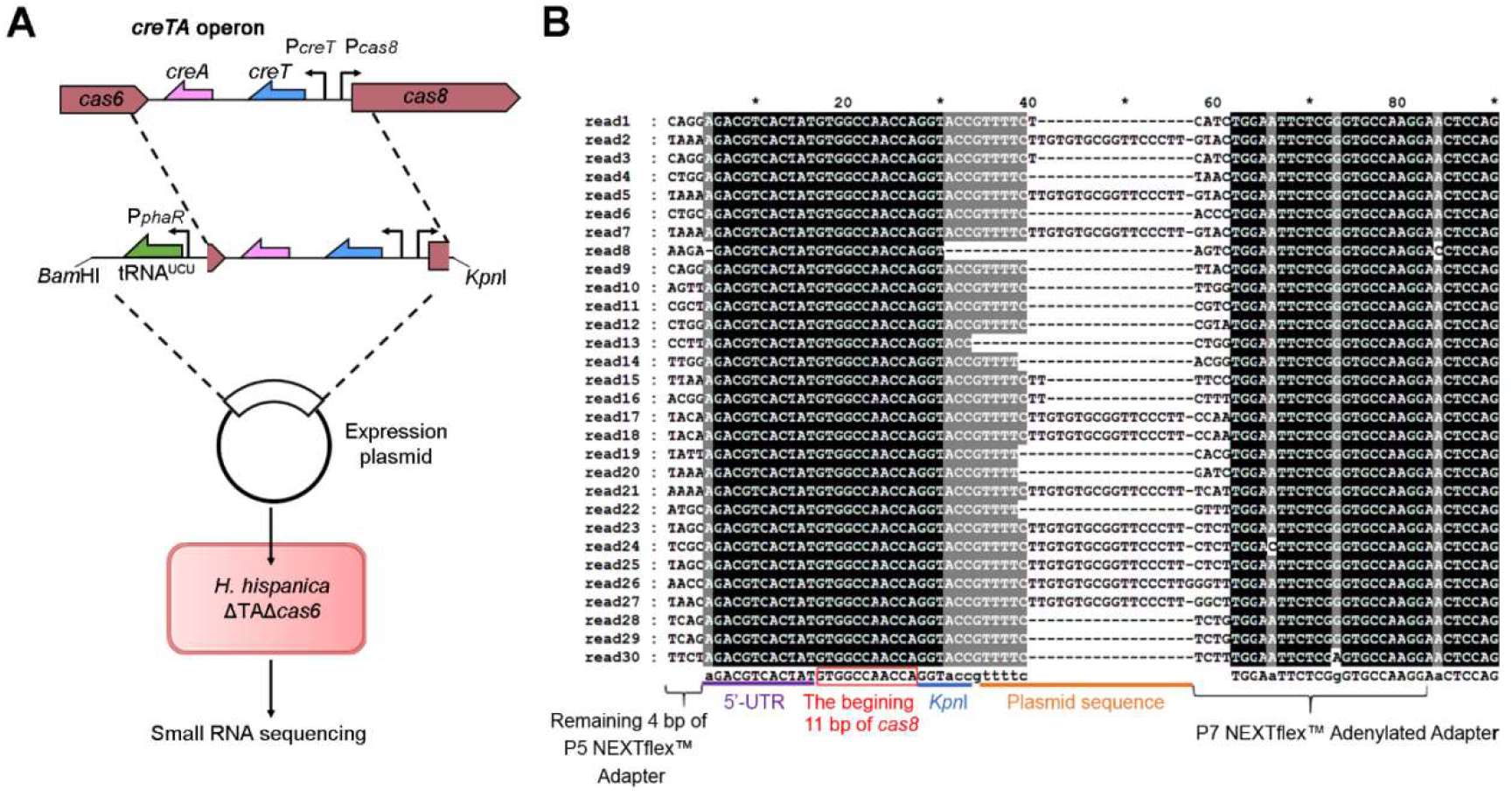
Determination of *cas8* TSS during a small RNA sequencing (sRNA-seq) assay, related to Figure 1. (**A**) The procedure of an sRNA-seq assay designed to analyze the transcription profile of *creTA* operon. The chromosomal region including *creTA* and flanking sequences from *cas6* and *cas8* were cloned into the expression plasmid pWL502. The recombinant plasmid was introduced into the *ΔTAΔcas6* cells of *H. hispanica* (so that *P_creT_* and *P_cas8_* were both de-repressed), from which the small RNA was extracted and subjected to sequencing. Note that the recombinant plasmid was designed to over-express tRNA^UCU^ to suppress the toxicity of CreT (Li *et al*., 2021). **(B)**Example reads revealing the TSS of *cas8*. Nucleotides corresponding to the 5’ untranslated region (5’-UTR) and the beginning 11 bp of *cas8* ORF are indicated.

**Figure S2.**
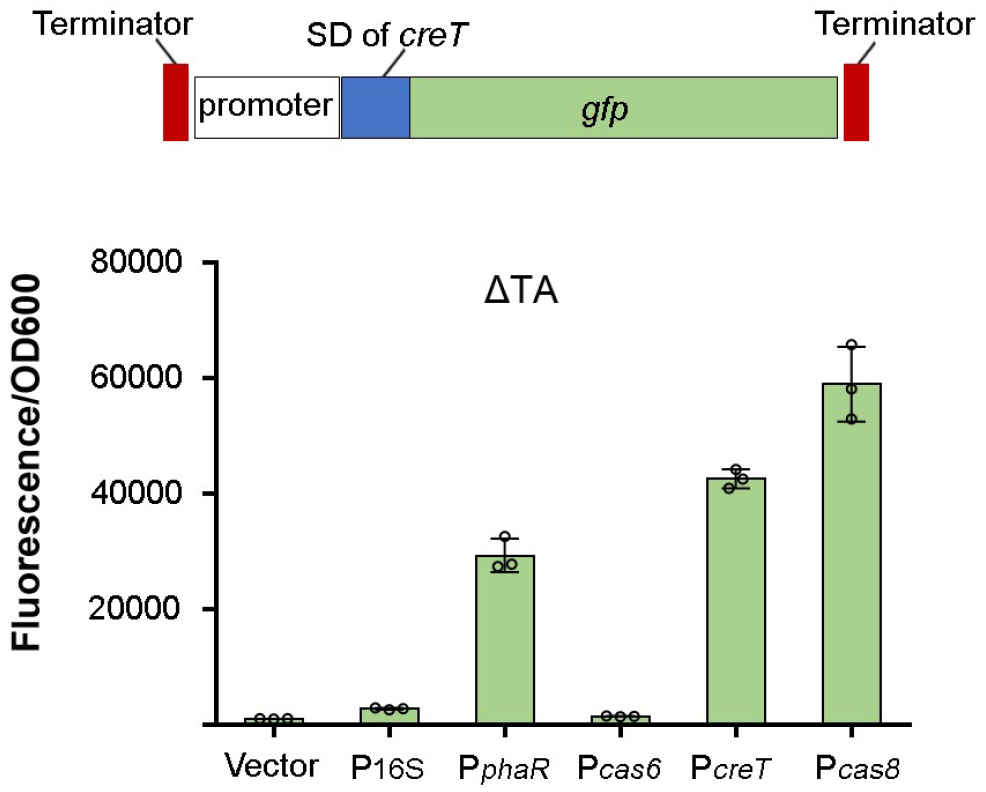
Probe the strength of *P_cas8_* using the *gfp* reporter gene, related to Figure 1. DNA constructs containing *gfp* controlled by different promoters were cloned into the expression plasmid pWL502 and then introduced into ΔTA cells (lacking the repressing CreA molecules) for fluorescence determination. To facilitate strength comparison among different promoters, an identical SD sequence (the one of *creT*) was employed. P16S, the promoter of the 16S rRNA gene. P_*phaR*_, a strong constitutive promoter conventionally used for gene over-expression in haloarchaea (Cai *et al*., 2015). Terminator, a string of eight thymines. Error bars, mean ± s. d. (n =3).

**Figure S3.**
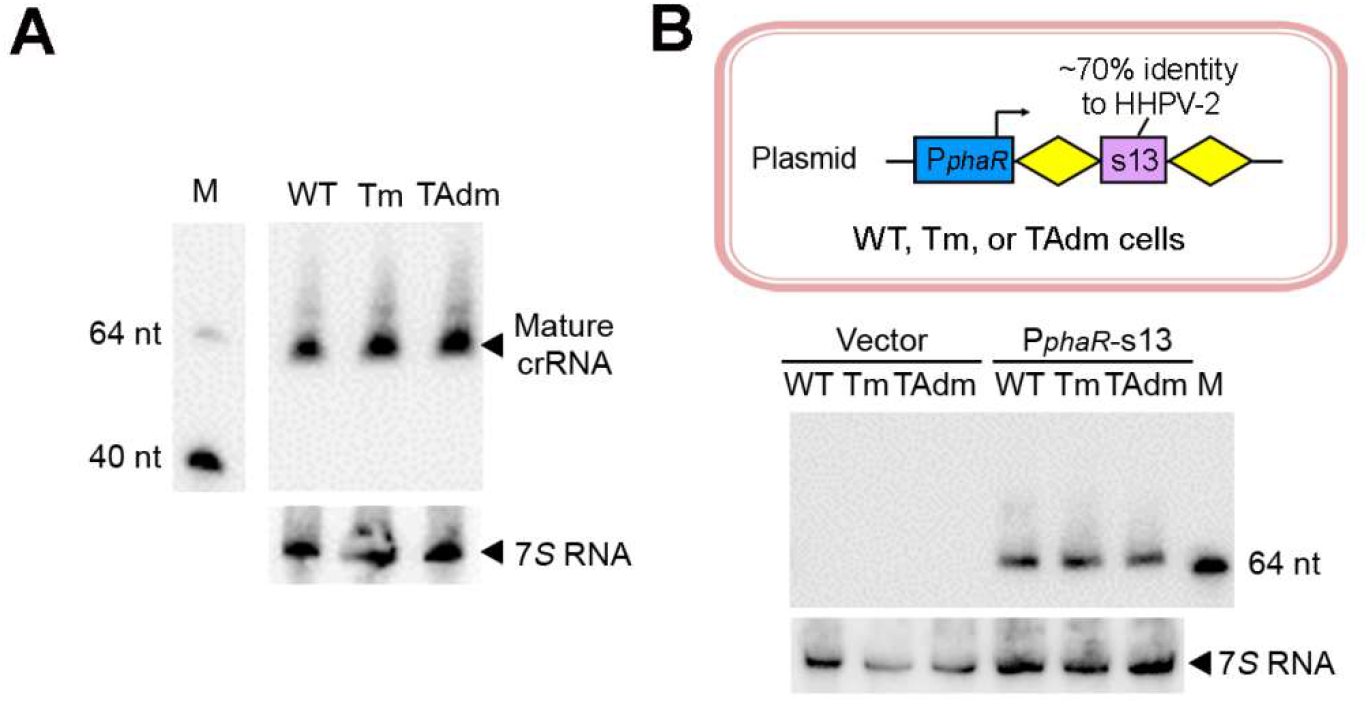
Northern blotting of total crRNA (A) or the crRNA of s13 (B) in WT, Tm and TAdm cells, related to Figure 2. In panel B, a plasmid over-expressing the crRNA of s13 spacer was introduced into the three hosts. Note that, in the CRISPR array on *H. hispanica* chromosome, the s13 spacer (the teriminal one) is followed by a degenerated repeat and its crRNA products could hardly be detected by Northern blotting (consistent to our previous observation (Gong *et al*., 2019)), while the s13 spacer on the plasmid was designed between two typical repeats (represented by two yellow diamonds). 7*S* RNA served as the internal control. M, biotin-labeled oligonucleotides.

**Figure S4.**
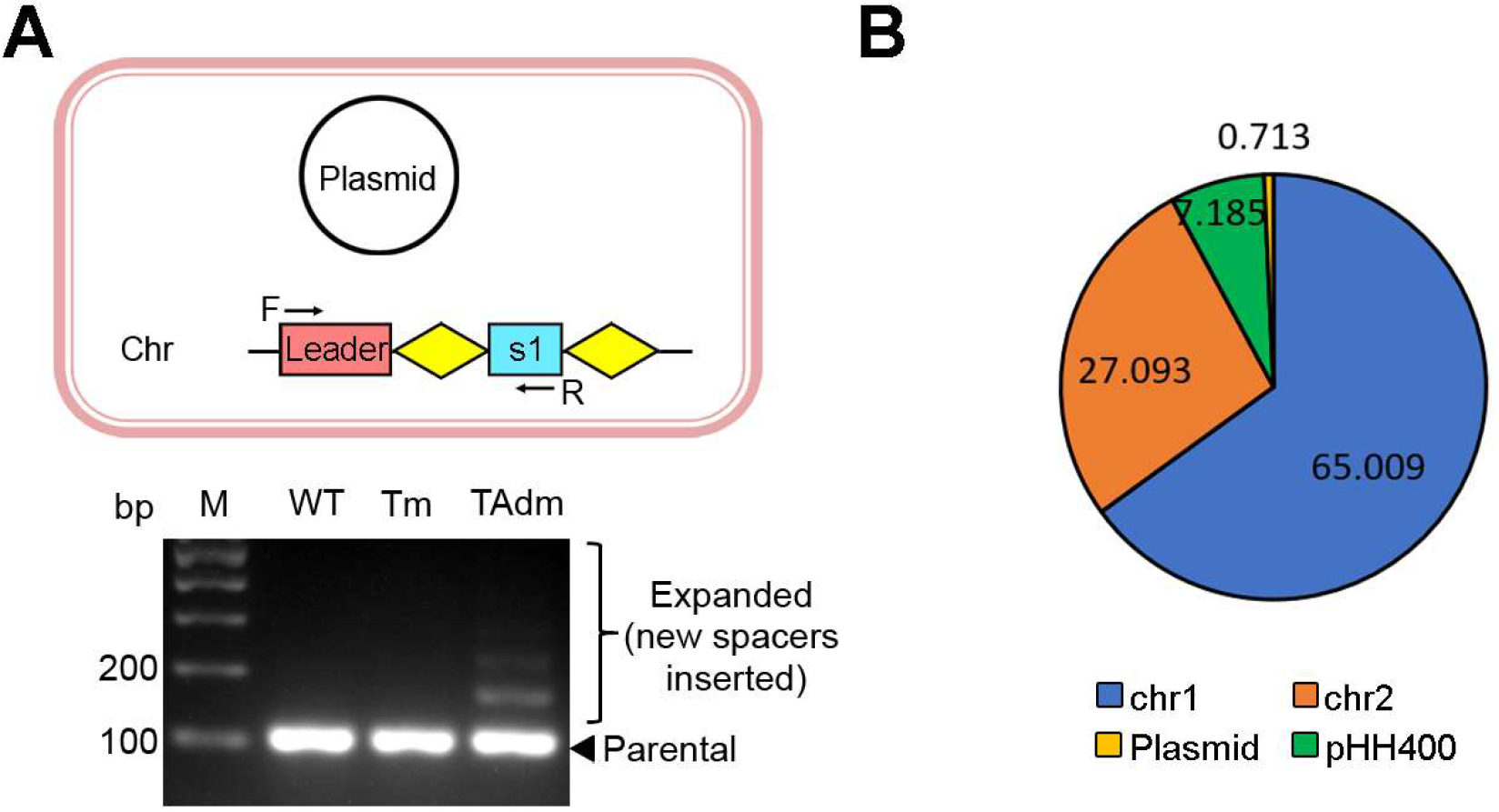
Naïve adaptation in TAdm cells, related to Figure 2. **(A)** *H. hispanica* cells containing the empty vector pWL502 were cultivated for a long period and spacer acquisition was monitored by PCR analysis. Two primers against the leader sequence and the first CRISPR spacer (s1), respectively, were used for amplification. **(B)**Scheme showing the derivation ratio (%) of new spacers, which was analyzed by illumine sequencing.

**Figure S5.**
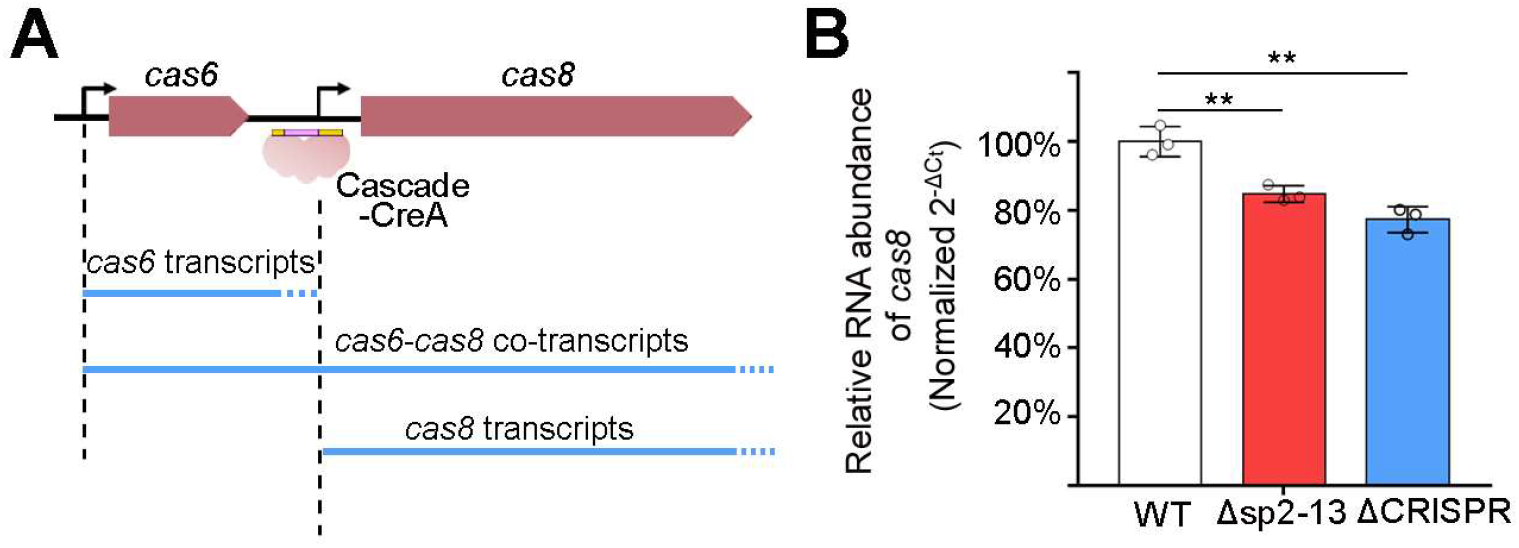
The RNA level of *cas8* relative to *cas6* in *H. hispanica* cells that contain different number of CRISPR spacers, related to Figure 3. **(A)**Scheme depicting the gene organization of *cas6* and *cas8* and their potential transcripts. CreA directs Cascade to bind to the DNA of P*cas8*, which will suppress its activity and in principle also attenuate the readthrough transcripts driven by *P_cas6_*. **(B)**The qPCR assay to determine the relative RNA abundance of *cas8* in WT, Δsp2-13 or ΔCRISPR cells. The Δsp2-13 mutant contains only one spacer. RNA of *cas6* served as the internal control. Error bars, mean±s.d. (n=3); two-tailed Student’s *t* test [**P < 0. 01].

**Figure S6.**
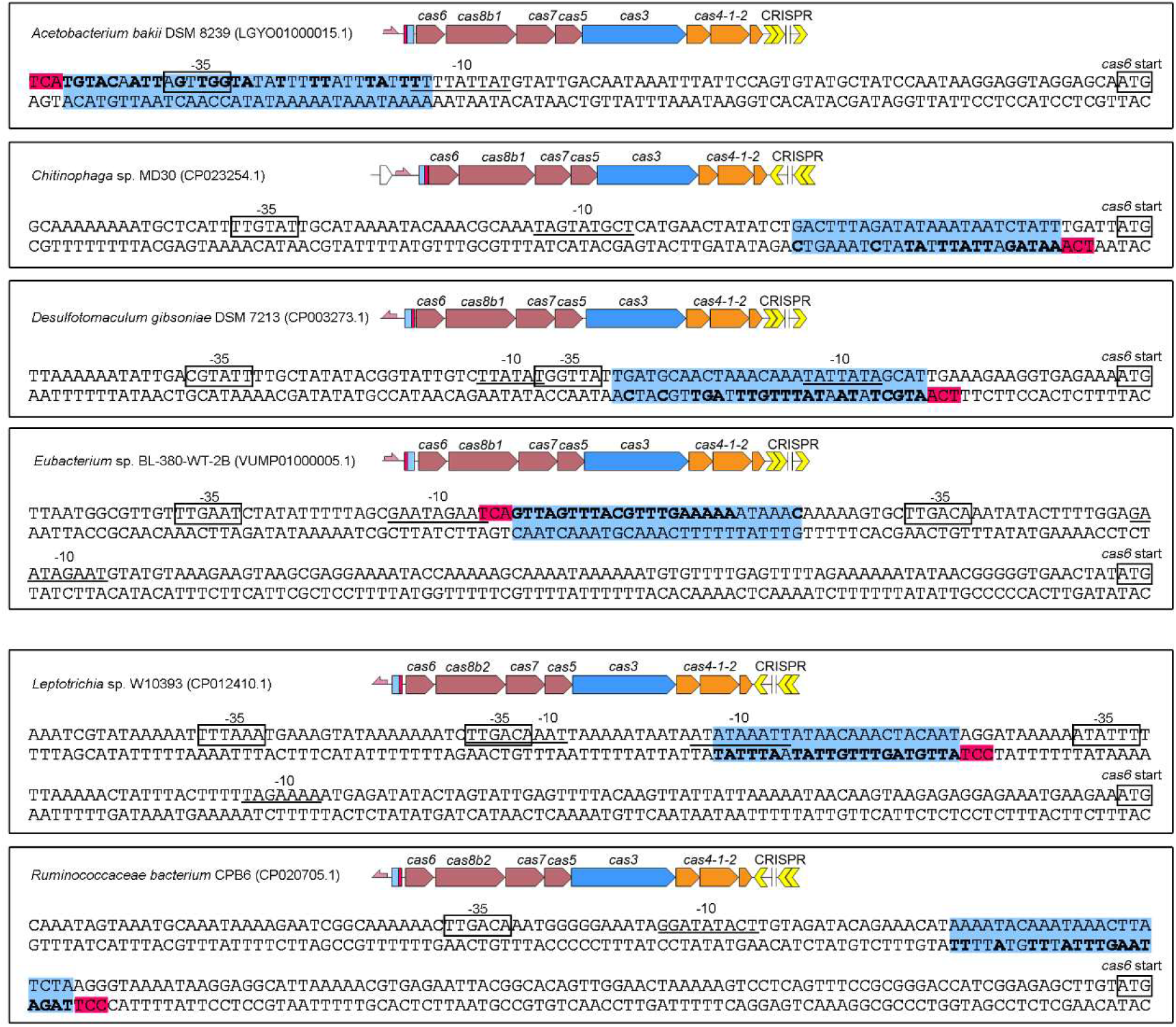
The proximity of CreR (or CreA) target site to the *cas* promoter in some type I-B systems, related to Figure 4. The target sequence (protospacer) of CreR and the PAM (protospacer adjacent motif) nucleotides are indicated with blue and red background colors, respectively. Nucleotides in bold indicate the identical ones shared between the ‘spacer’ (ΨS) of CreR and its corresponding protospacer. The −35 and −10 promoter elements were predicted using the Softberry web server (http://www.softberry.com/).

**Figure S7.**
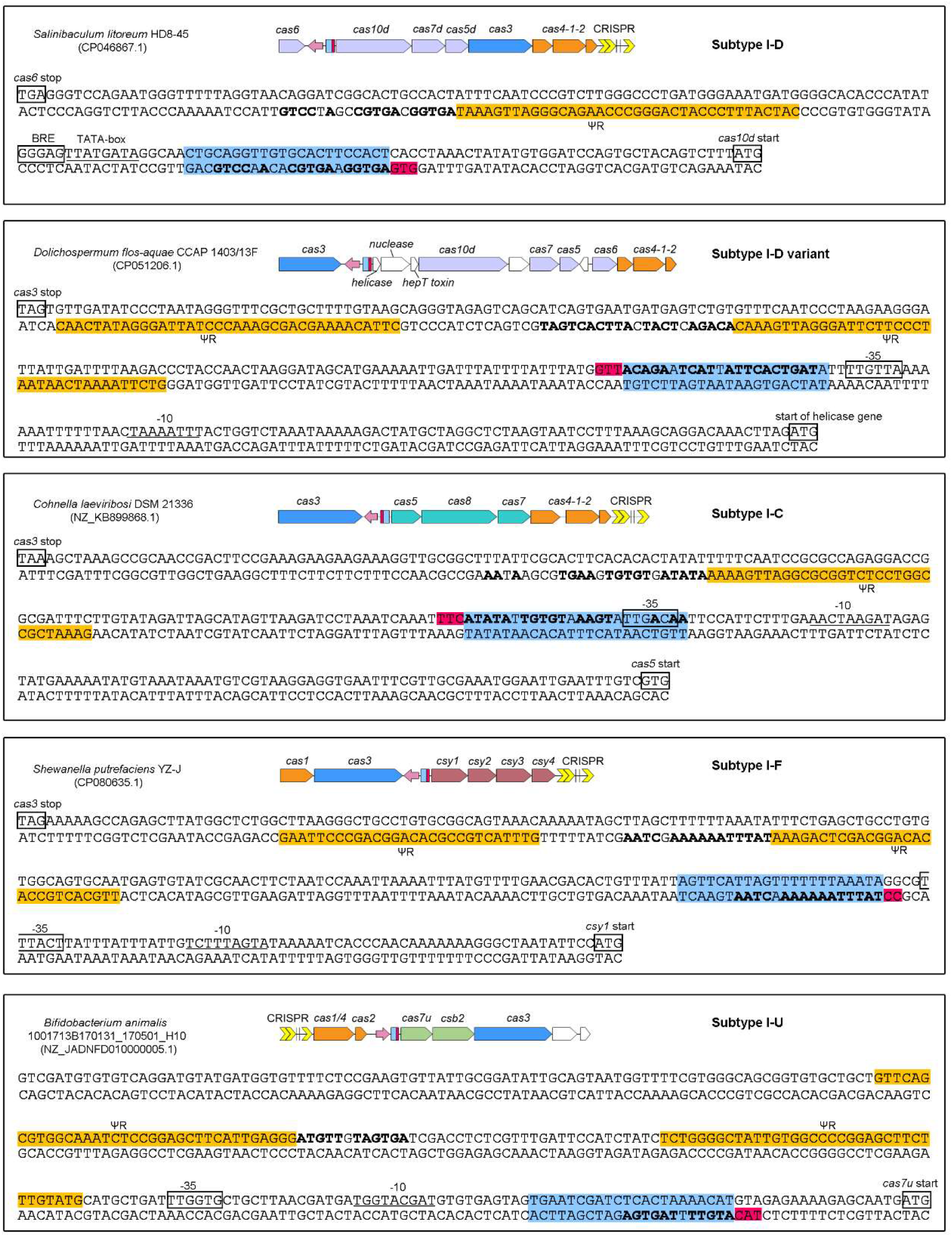
The proximity of CreR target site to the promoter of CRISPR effector genes in different Class1 subtypes, related to Figure 4. The target sequence (protospacer) of CreR and the PAM (protospacer adjacent motif) nucleotides are indicated with blue and red background colors, respectively. Nucleotides in bold indicate the identical ones shared between the ‘ spacer’ (ΨS) of *creR* and its corresponding protospacer. The −35 and −10 promoter elements were predicted using the Softberry web server (http://www.softberry.com/).

**Figure S8.**
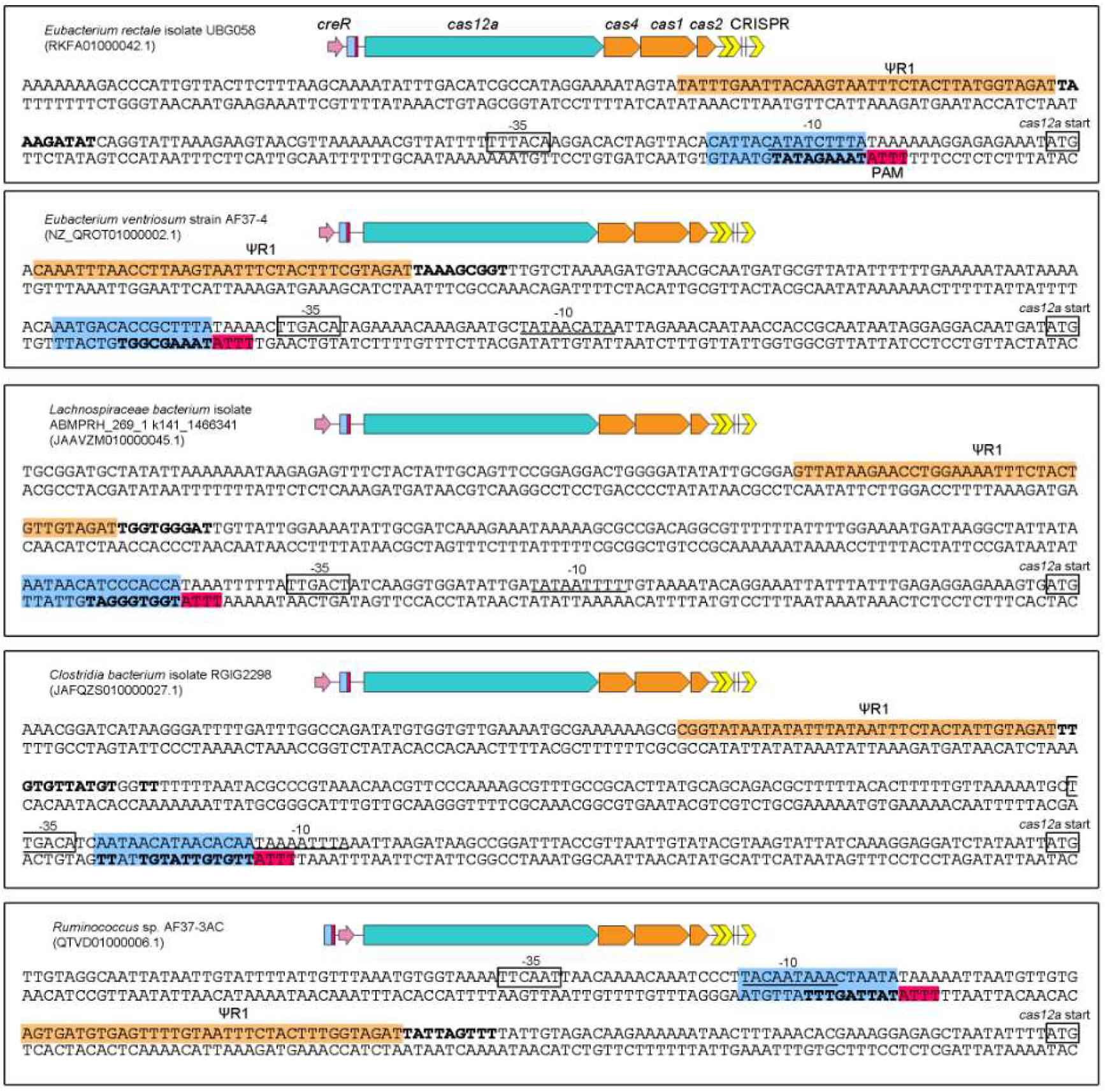
The proximity of CreR target site to the promoter of V-A CRISPR effector gene (*cas12a*), related to Figure 5. The target sequence (protospacer) of CreR and the PAM (protospacer adjacent motif) nucleotides are indicated with blue and red background colors, respectively. Nucleotides in bold indicate the identical ones shared between the ‘ spacer’ (ΨS) of *creR* and its corresponding protospacer. ΨR2 was difficult to predict for its extensive degeneration. The −35 and −10 promoter elements were predicted using the Softberry web server (http://www.softberry.com/).

**Figure S9.**
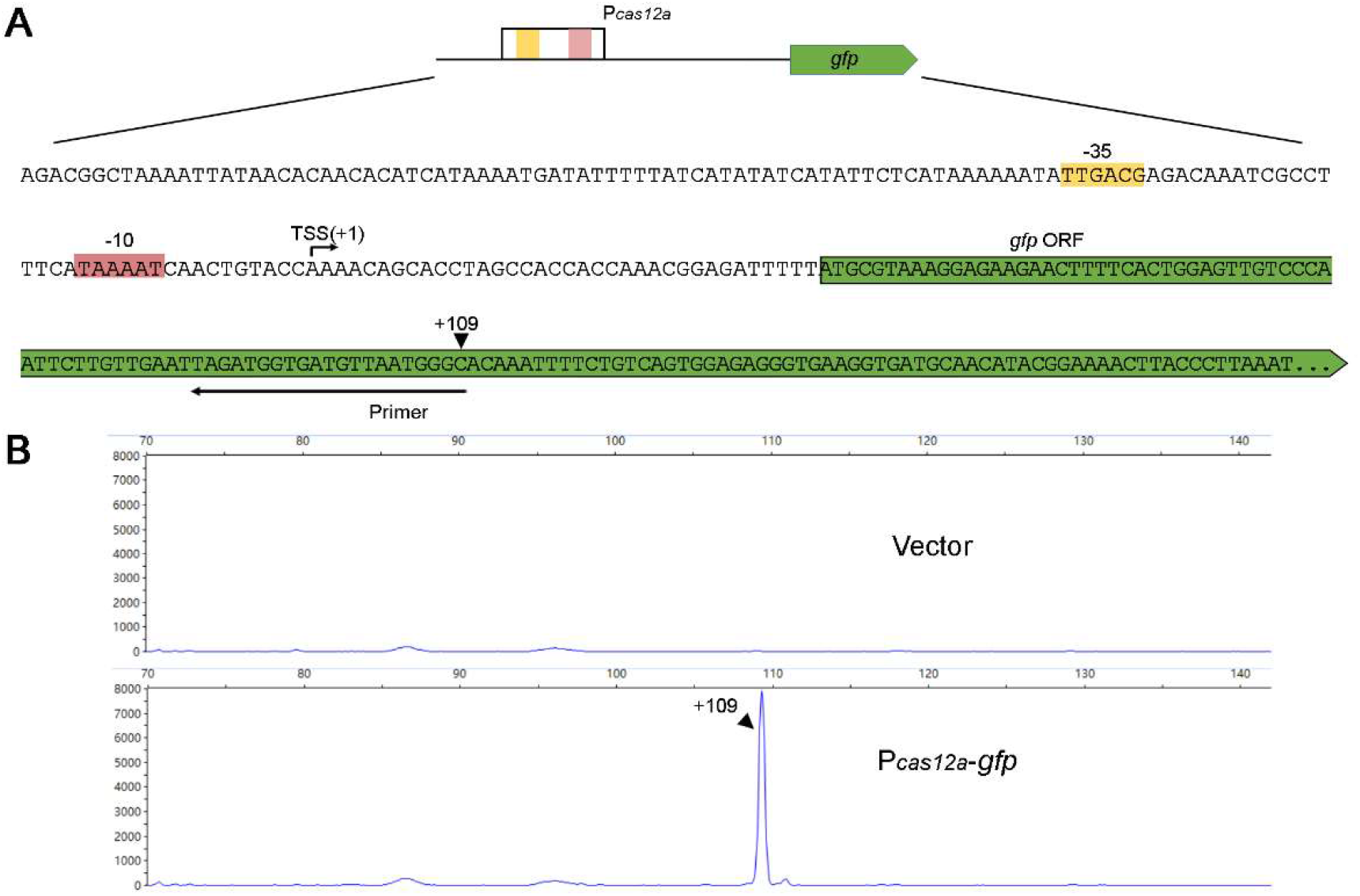
Primer extension assay to determine the transcription start site (TSS) of the *cas12a* promoter, related to Figure 4. **(A)**Scheme showing the experimental design. The modified pACYC plasmid carrying the *P_cas12a_-gfp* construct was introduced into *E. coli* cells, and the total RNA was extracted for primer extension. The primer was designed against the *gfp* RNA transcript and 5’-labeled by FAM (see the Methods part). The cDNA products of primer extension assay were subjected to fragment size analysis. **(B)**The results of primer extension. TSS was identified according to the fragment size of the cDNA products. Cells containing the empty vector were used as the negative control.

